# Lipid droplet dysmetabolism affects cell homeostasis in an *in vitro* model of Alzheimer’s disease

**DOI:** 10.1101/2025.06.09.658644

**Authors:** Tânia Fernandes, Margarida Caldeira, Tânia Melo, Bruna B. Neves, Cristina Carvalho, Rosa Resende, Riccardo Filadi, Paola Pizzo, M. Rosário Domingues, Cláudia F. Pereira, Paula I. Moreira

**Author notes:** **Corresponding authors:** Tânia Fernandes, Cláudia F. Pereira, and Paula I. Moreira, Center for Neuroscience and Cell Biology, University of Coimbra, Rua Larga, Faculty of Medicine, Polo I, 1st floor, 3004-504 Coimbra, Portugal. (Tânia Fernandes), (Cláudia F. Pereira), and (Paula I. Moreira). Authors contributed equally to this work (Cláudia F. Pereira, Paula I. Moreira).

## Abstract

Lipid droplets (LD) are dynamic organelles involved in neutral lipid storage, energy homeostasis, and can prevent lipotoxicity and oxidative distress. LD dysmetabolism has been considered a pathological hallmark in neurodegenerative disorders, including Alzheimer’s disease (AD). In this study, we investigated the alterations in LD metabolism and their impact on mitochondria-associated membranes (MAM) in an in vitro model of AD, namely the mouse neuroblastoma cell (N2A) line overexpressing the amyloid precursor protein with the familial Swedish mutation (APPswe). In APPswe cells, we found depletion of LD associated with an accumulation of free fatty acids that can be related with the observed LD degradation by chaperone-mediated autophagy. In these cells we also found decreased levels of seipin, which might contribute to triacylglycerol accumulation. These lipid alterations are associated with increased levels of ROS and lipid peroxidation in APPswe cells. The pharmacological modulation of DGAT1, that mediates triacylglycerol synthesis, normalized LD size and improved ER-mitochondria contacts and mitochondrial function in APPswe cells. In summary, these observations suggest the involvement of altered LD metabolism in AD pathophysiology, which impact on MAM and mitochondria function leading to cell dyshomeostasis. Our findings also support the idea that LD are relevant therapeutic targets in AD.

**Graphical abstract:** 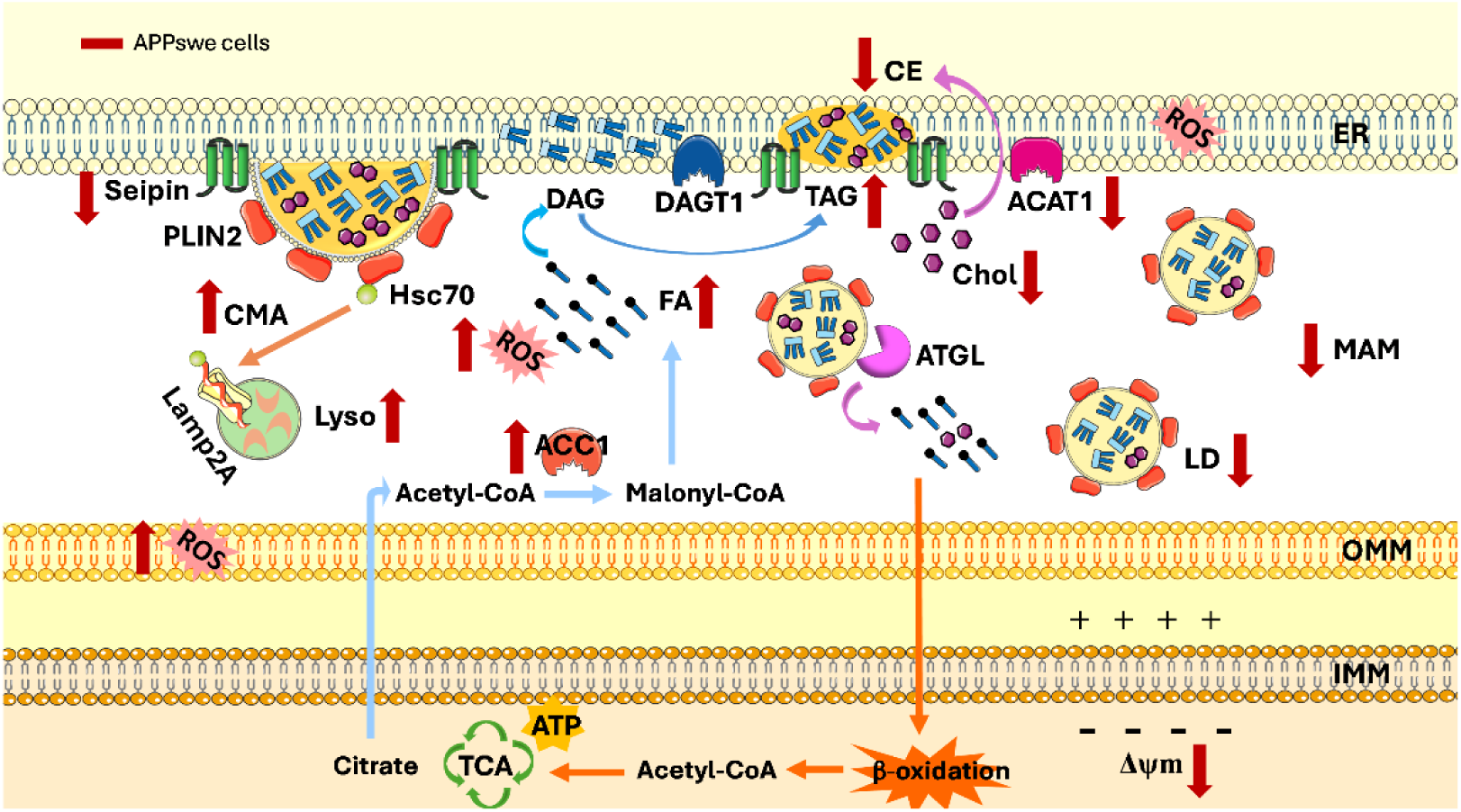

## 1. Introduction

Lipid droplet (LD) is a neutral lipid reservoir that can provide substrates for membrane formation and energy metabolism and can also prevent lipotoxicity and oxidative distress [1–3]. These organelles are composed by a phospholipid monolayer with integral- and peripheral-associated regulatory proteins, such as perilipin (PLIN), that surround a neutral lipid core formed by triacylglycerol (TAG) and cholesteryl ester (CE) [2,4,5]. The LD biogenesis begins in the endoplasmic reticulum (ER) with the esterification of activated fatty acid (FA) to diacylglycerol (DAG) or a sterol (such as cholesterol) that are then converted to TAG and CE mediated by DAG acyltransferases (DGAT1 and 2) and acyl-CoA:cholesterol O-acyltransferases (ACAT/SOAT), respectively [2,6,7]. The accumulation of neutral lipids between ER leaflets allows the formation of nascent LD [8]. Several factors involved in LD biogenesis, including seipin, a protein found in the ER membrane- and mitochondria-associated membrane (MAM), are recruited to the lens structure allowing the growth of the nascent LD [9–11].

LD interact with other organelles, including the ER, mitochondria, and lysosomes to promote the inter-organelle transfer of lipids upon the degradation of LD to release FA from stored TAG to be used for energy production, and to decrease the cytosolic levels of free fatty acid (FFA) avoiding lipotoxicity and/or aberrant lipid signaling [7,12,13]. The interaction between LD and lysosomes facilitates the degradation of the LD core, as well as the degradation of PLIN that surrounds the LD membrane, allowing the mobilization of its content for energy production to sustain metabolic processes and membrane biosynthesis [2]. LD can be degraded by several pathways, including macroautophagy and chaperone-mediated autophagy (CMA). In macroautophagy, portions of LD can be engulfed by the phagophore membrane and the autophagosome fuses with the lysosome for degradation of neutral lipids [2,14]. In CMA, proteins with specific amino acids characteristics (KFERQ motif), such as PLIN2 and PLIN3, are recognized by the cytoplasmic heat shock-associated protein 70 (hsc70) and transported to the lysosomal membrane where they interact with the lysosome-associated membrane receptor type 2A (Lamp2A) being internalized into the lysosome [15].

Although LD biology has been well studied in hepatic [16,17] and cardiovascular [18] disorders, the role of these organelles in aging and several other pathological conditions are now gaining momentum [2,19]. In fact, brain lipid dysmetabolism plays an important role in the pathogenesis and progression of several neurodegenerative disorders, including Alzheimer’s disease (AD) [20–22].

AD is an age-related neurodegenerative disorder characterized by the deposition of extracellular amyloid β (Aβ) peptide and intracellular hyperphosphorylated tau protein, and progressive neuronal loss leading to cognitive impairment [23,24]. Other pathophysiological alterations in AD are altered MAM structure and function, including changes in MAM-associated lipid metabolism and Ca^2+^ signaling, increased ER stress, and mitochondrial dysfunction [25–27]. Alterations in LD metabolism have been reported in several AD models, including an accumulation of LD in fibroblasts from familial and sporadic AD (FAD and SAD, respectively) patients and in presenilin-1 (PSEN1)-mutant mouse embryonic fibroblasts (MEFs) and in PSEN1 knockdown (PS1-KD) cells [28]. In PSEN1 knockout (KO) MEFs an increase of cholesteryl esterification and LD was also observed [28]. It has been also shown that increased reactive oxygen species (ROS) levels and the glia-neurons lactate shuttle contribute to lipid synthesis in neurons and to LD accumulation in glia via apolipoprotein E/D (APOE/D) [29]. The increase of lipids stored in LD was also reported in peripheral blood plasma and in the frontal cortex of AD patients [30–32] and in PSEN1/amyloid precursor protein (APP) and APP/tau mice brains [32,33]. Recent studies performed in transgenic mice and differentiated induced pluripotent stem cells highlight the microglial damage of LD in AD [34,35]. However, the alteration of LD metabolism in AD has not been explored.

The clarification of the mechanisms underlying lipid dysregulation in AD is crucial to identifying reliable biomarkers and therapeutic targets for the diagnosis and development of effective strategies to fight this devastating neurodegenerative disease. In this line, this study was aimed at investigating the alterations in LD metabolism in a neuroblastoma cell line and their effects in MAM, mitochondrial function and cellular homeostasis using an *in vitro* model of AD, namely the mouse neuroblastoma cell line (N2A) overexpressing APP containing the familial Swedish mutation (APPswe).

## 2. Materials and methods

### 2.1. Cell culture and treatments

The wild type (WT) mouse neuroblastoma cell line (N2AWT) and the N2A APPswe cells that stably overexpress the human Swedish mutant APP KM670/671NL [36], were cultured in Dulbecco’s Modified Eagle’s medium (DMEM, Sigma-Aldrich, D5648) supplemented with 10% (*v/v*) fetal bovine serum (FBS) (Gibco, 26400-044), 3.7 g/L sodium bicarbonate (Merck, S8761), 1% (*v/v*) non-essential amino acids (Merck, M7145), and 1 mM sodium pyruvate (Merck, S8636), as previously described [25,37]. Cell culture medium was supplemented with 1% (*v/v*) penicillin/streptomycin (Gibco, 15140-122) or 0.4 mg/mL geneticin (Gibco, 10131027) for WT or APPswe cells, respectively. To induce starvation, cells were maintained in the same cell culture medium supplemented with 1% (*v/v*) FBS. Cells were cultured at 37°C in a humidified 5% CO2-95% air atmosphere.

For the ubiquitin-proteosome system (UPS) study, cells were treated with 50 nM MG132 (Calbiochem, 474790), a proteosome inhibitor, during 24 h. To stimulate LD formation, cells were incubated with 25 µM oleic acid (OA) (Thermo Scientific, 31997) during 24 h, a well know inducer of LD accumulation through its esterification to glycerol by DGAT [38]. Lastly, for the inhibition of LD formation cells were treated with 20 µM of A922500 (Sigma, A1737), an inhibitor of DGAT1, during 48 h. In all the previous treatments, cells were maintained in medium supplemented with 10% (*v/v*) FBS. To study macroautophagy, cells were treated with 25 µM OA, during 24 h, washed 1x with phosphate-buffered saline (PBS) to remove the OA and incubated during 48 h with 10 µM chloroquine (CQ) (Sigma, C6628), an inhibitor of autophagy, under starvation (medium supplemented with 1% (*v/v*) FBS). Lastly, for the CMA assay, cells were treated with 25 µM OA, during 24 h, washed 1x with PBS to remove the OA, and incubated with 10% (*v/v*) FBS or 1% (*v/v*) FBS (starvation) medium during 24 h. Cells were maintained at 37°C in a humidified 5% CO_2_-95% air atmosphere.

### 2.2. Cell protein extraction

To evaluate protein levels, cells were detached by trypsinization and centrifuged at 200 x g for 5 min, at 4°C. Homogenization of pellets was gently performed with a glass/teflon homogenizer in isolation buffer, pH 7.4, composed of 225 mM mannitol, 75 mM sucrose, 30 mM Tris-HCl and 0.1 mM EGTA supplemented with a 1% (*v/v*) cocktail of proteases inhibitors (Sigma, P2714). The suspension was centrifuged 8x at 600 x g, for 5 min, at 4°C, to completely remove cell debris and nuclei and the supernatant was collected.

To analyze microtubule-associated protein light chain 3 (LC3) levels, cells were cultured in 6-well plate (187,500 cells per well) and, after 24 h of incubation, they were treated with 20 µM CQ during 24 h, at 37°C, in a humidified 5% CO2-95% air atmosphere. Cells without CQ treatment were used as control cells. After the incubation period, cells were lysed by adding 40 µl RIPA buffer (250 mM NaCl, 50 mM Tris, 1% Nonidet P-40, 0.5% DOC, and 0.1% SDS, pH 8.0) supplemented with proteases inhibitors (Sigma, P2714), 0.2 mM phenylmethanesulfonyl fluoride (PMSF) (Sigma, P7626), 5 mM sodium fluoride (NaF), 1 mM sodium ortovanadate, 1 mM dithiothreitol (DTT) (Amresco, 281), and 1% Triton X-100. The mixture was incubated during 20 min in ice to favor cells disruption, and then centrifugated, at 4°C, during 10 min, at 18,000 x g (Eppendorf centrifuge 5415C). The supernatants, which represent the cytosolic fraction, were collected.

The protein concentration was measured using the Pierce bicinchoninic acid (BCA) Protein Assay Kit (Thermo Fisher, 23227).

### 2.3. Western blot analysis of proteins

The samples were boiled in loading buffer for denaturation. Sample proteins (60 µg) were separated using 5-12% SDS-PAGE, transferred to a polyvinylidene difluoride (PVDF) membrane (Millipore), and blocked in 5% (*w/v*) BSA prepared in Tris-buffered saline containing 0.1% (*v/v*) Tween-20 (TBS-T). Then, the membranes were incubated overnight at 4°C with the primary antibody, washed with TBS-T, and incubated at room temperature (RT) for 1 h with the secondary antibody. Membranes were developed in ChemiDoc Imaging System (Bio-Rad, Hercules, California, USA) using enhanced chemifluorescence (ECF) substrate (GE Healthcare, RPN5785), and quantifications performed in the Bio-Rad Image Lab Software 6.1. Primary and secondary antibodies utilized in WB assays are summarized in Table 1. Actin protein was used as loading control for all protein analyzed, except for ACC1 protein that was used Ponceau S staining.

**Table 1.** Primary and secondary antibodies utilized for Western blot (WB), immunofluorescence (IF) staining and proximity ligation assay (PLA) analyses.

| Antibody | Dilution | Species | Company | Catalog N°. |
| --- | --- | --- | --- | --- |
| β-Actin | 1:10,000 | Mouse | Sigma-Aldrich | A8717 |
| ACC | 1:1,000 | Rabbit | Invitrogen | MA5-15025 |
| ADRP/Perilipin2<br>(PLIN2) | 1:250 | Rabbit | Proteintech | 15294-1-AP |
| DGAT1 | 1:2,000 | Goat | Abcam | ab59034 |
| IP3R1 | 1:1,000 | Rabbit | Abcam | ab5804 |
| Lamp1 | 1:100 | Mouse | Santa Cruz<br>Biotechnology | sc-20011 |
| Lamp2A | 1:100 | Rat | DSHB | ABL-93 |
| LC3B | 1:1000 (WB)<br>1:300 (IF) | Rabbit | Cell Signalling | 3868S |
| Seipin | 1:500 | Rabbit | Invitrogen | PA5-114918 |
| SOAT1/ACAT1 | 1:500 | Rabbit | Abcam | ab307597 |
| TOM20 | 1:200 | Rabbit | Santa Cruz<br>Biotechnology | sc-11415 |
| VDAC1 | 1:250 | Mouse | Abcam | ab14734 |
| Anti-goat IgG | 1:10,000 | Rabbit | Santa Cruz<br>Biotechnology | sc-2771 |
| Anti-mouse IgG | 1:10,000 | Goat | Thermo Fisher | 31320 |
| Ati-Rabbit IgG | 1:20,000 | Goat | GE Healthcare | NIF1317 |
| Anti-Mouse<br>IgG Alexa Fluor 488 | 1:200 | Goat | Invitrogen | A11001 |
| Anti-Rabbit<br>IgG Alexa Fluor 488 | 1:200 | Goat | Invitrogen | A11008 |
| Anti-Rat<br>IgG Alexa Fluor 594 | 1:500 | Goat | Invitrogen | A11007 |

### 2.4. Analysis of LD turnover by immunofluorescence and confocal microscopy assays

For confocal imaging, WT and APPswe cells were cultured in Ibi-treated µ-Slide 8 well chamber (Ibidi, 80806) (20,000 cells per well) and incubated for 72 h in a humidified 5% CO_2_-95% air atmosphere at 37°C.

#### LD staining

After treatments, cells were washed 2x with warm PBS, fixed with 4% (*w/v*) paraformaldehyde (PFA) (Thermo Scientific, A11313) in PBS during 15 min at RT, washed 3x with PBS, and incubated with 1:200 LipidTox Red (Invitrogen, H34476) in PBS for 2 h at RT. After washing 2x with PBS, cells were incubated with 15 µg/mL Hoechst 33342 (Molecular Probes) for 10 min to stain nuclei. Cells were washed 2x with PBS and imaging was immediately performed.

#### LD interaction with lysosomes

Cells were washed 2x with warm PBS, fixed with 4% (*w/v*) PFA in PBS during 15 min at RT, and washed again 3x with PBS. Cells were permeabilized with 0.1% (*w/v*) Triton X-100 (Sigma, T8787) in PBS for 5 min at RT, washed 3x with PBS, and blocked for 1 h at RT in blocking solution [3% (*w/v*) BSA and 0.05% (*w/v*) saponin (Alfa Aesar, A18820) in PBS]. Then, cells were incubated overnight at 4°C, with anti-Lamp1 antibody diluted in the blocking solution. Cells were then washed 3x in PBS and incubated for 1h at RT, with the secondary antibody Alexa Fluor 488 diluted in the blocking solution. Lastly, cells were washed 3x in PBS and incubated with LipidTox Red and Hoechst 33342 (Molecular Probes), as described above.

#### CMA-mediated LD degradation and LD interaction with LC3

After treatments, cells were washed 2x with warm PBS, fixed with 4% (*w/v*) PFA in PBS during 15 min at RT, and washed 3x with PBS. Cells were permeabilized with 0.1% (*v/v*) Triton in PBS for 5 min at RT, washed 3x with PBS, and blocked during 30 min at 37°C in PBS containing 10% (w/v) BSA (Nzytech, 9048). Then, cells were incubated overnight at 4°C, with anti-LC3B or with anti-ADRP/Perilipin 2 (PLIN2), and anti-Lamp2A diluted in the incubation solution [PBS with 3% (*w/v*) BSA]. Cells were then washed 6x with PBS for 2 min each wash and incubated for 45 min at 37°C with the secondary antibodies, Alexa Fluor 488 and Alexa Fluor 594 diluted in the incubation solution. Lastly, cells were washed 6x with PBS for 2 min each wash and, for LC3-LD interaction, cells were incubated with LipidTox Red and Hoechst 33342, as described above, and washed 2x with PBS before the addition of Aqua-Poly/mount (Polysciences, 18606). Imaging of LC3-LD interaction was immediately performed.

All the images were acquired with a Zeiss LSM 710 Confocal Microscope (Carl Zeiss, Jena, Germany) with a Plan-Apochromat 63×/1.4 oil objective (Carl Zeiss) and with the following laser lines: diode 405 nm, argon/2 488 nm, and DPSS 561 nm. Three-dimensional imaging of the cells was performed using the Z-stack function with an interval of 0.57 µm ranging the entire thickness of the cells. For image colocalization studies the JACoP plugin of the Fiji software was used, generating the Mander’s correlation coefficient. The LD number was calculated using the Spot detection tool of Imaris software.

Primary and secondary antibodies used for immunofluorescence are summarized in Table 1.

### 2.5. Quantification of ER-mitochondria contacts by confocal microscopy

#### Proximity ligation assay (PLA)

Cells were cultured in Ibi-treated µ-Slide 8 well chamber (20,000 cells per well) and, after treatment with A922500, cells were washed 2x with PBS, fixed with 4% (*w/v*) PFA in PBS during 15 min at RT, and permeabilized with 1% (*v/v*) Triton in PBS for 5 min at RT. PLA was performed on the permeabilized cells by using the DuoLink FarRed In Situ Detection Kit (Sigma-Aldrich, DUO92013) and DuoLink anti-mouse (Sigma-Aldrich, DUO92002) and anti-rabbit (Sigma-Aldrich, DUO92004), following the manufacturers’ instructions. Cells were treated with the blocking solution [3% (w/v) BSA supplemented with 0.2% (*v/v*) Tween-20] during 1 h at RT, incubated with primary antibodies anti-IP3R1 and anti-VDAC1 (Table 1) overnight at 4°C, and then incubated with PLA probes at 37°C for 1 h. After washing 2x during 5 min, the ligation-ligase solution was added and incubated for 30 min at 37°C. Next, cells were incubated with amplification-polymerase solution, at 37°C for 100 min, in the dark. Finally, cells were washed, incubated with 15 µg/mL Hoechst 33342 for 10 min, at RT, and washed 3x with PBS before the addition of Aqua-Poly/mount. Imaging was performed on a Zeiss LSM 710 Confocal Microscope (Carl Zeiss, Jena, Germany) with a Plan-Apochromat 63×/1.4 oil objective with the following laser lines: diode 405 nm and HeNe633. Quantification of the number of PLA probe dots was performed using the Spot detection tool of the Imaris software.

#### ER-mitochondria RspA-splitFAST probe

Cells were seeded in 18 mm coverslips (60,000 cells per coverslip) and after 24 h were treated with A922500. The day before imaging, cells were transfected with short RspA-CFAST (targeted to OMM) and RspA-NFAST (targeted to ER membranes) plasmids, as previously described [39], using Lipofectamine^TM^ 2000 (Thermo Fisher, 11668027).

Immediately before the imaging experiments, cells were stained with 0.17 µM MitoTracker Deep Red (Invitrogen, M22426), for 10 min in mKRB (140 mM NaCl, 2.8 mM KCl, 2 mM MgCl_2_, 1 mM CaCl_2_, 10 mM HEPES, and 10 mM glucose, pH 7.4, at 37°C). After washing with mKRB solution, the coverslips were mounted on a chamber and bathed in mKRB supplemented with 3 µM fluorogen HMBR (^TF^Lime) (Twinkle Factory, 480541-250).

The ER-mitochondria contact imaging was done using splitFAST-based probes visualized with a Leica TCS-II SP5 microscope, equipped with a Plan Apochromat 100x/1.4 N.A. objective and a tunable WLL laser. The WLL was set at 488 nm and 647 nm to excite the ER-mitochondria RspA-splitFAST probe and MitoTracker Deep Red, respectively. To visualize spatial overlaps between the ER-mitochondria RspA-splitFAST and MitoTracker Deep Red probes, the background was subtracted using ImageJ for both channels and preprocessed using a custom-written Python script. The Python libraries employed were openCV and scikit-mage. As previously described [39], the images were min-max normalized and filtered with a standard deviation of 1 using the “filters.gaussian” function from scikit-images. Images were binarized using a threshold set at half of the mean value of the pixel intensity values exceeding 20% of the pixel intensity distribution. The binarized images of ER-mitochondria RspA-splitFAST and MitoTracker were merged using pixel multiplication and the overlap fraction calculated on the MitoTracker image.

### 2.6. Quantification of long-chain free fatty acids and cholesterol levels

Cells were plated (1.25x10^6^ cells per T75) and incubated for 72 h in cell culture medium supplemented with 10% (*v/v*) FBS, at 37°C, in a humidified 5% CO_2_-95% air atmosphere. After, 8x10^6^ cells were collected and used to measure FFA and cholesterol levels. The concentration of long-chain FFA, total cholesterol and cholesteryl esters were determined by the Free Fatty Acid Quantitation Kit (Sigma-Aldrich, MAK044) and Cholesterol Quantitation Kit (Sigma-Aldrich; MAK043) following the manufacturers’ instructions.

### 2.7. Fluorimetric measurement of cytosolic and mitochondrial reactive oxygen species

Cytosolic ROS levels were measured at basal conditions with dichlorodihydrofluorescein diacetate (DCFH_2_-DA) (Life Technologies, C6827) and the levels of mitochondrial hydrogen peroxide (H_2_O_2_) were determined using the selective mitochondrial fluorescent probe MitoPY1 (Tocris Bioscience^TM^, 44-281-0) [40]. Cells were seeded in 48-well plate (18,750 cells per well), in triplicate, and incubated at 37°C, in a humidified 5% CO_2_-95% air atmosphere, for 72 h. For cytosolic ROS levels, cells were washed 1x with warm PBS to remove the culture medium and incubated for 30 min with 5 µM DCFH_2_-DA in Krebs solution (140 mM NaCl, 5 mM KCl, 1.5 mM CaCl_2_, 1 mM NaH_2_PO_4_, 9.6 mM glucose, 20 mM Hepes, 1 mM MgCl_2_, pH 7.4), at 37°C, in a humidified 5% CO_2_-95% air atmosphere. The incubation solution was replaced by fresh Krebs solution and the fluorescence signal was measured at 37°C for 60 min (λEx = 480 nm and λEm = 550 nm) using the microplate reader spectraMax iD3 (Molecular Devices, San Jose, CA, USA). For H_2_O_2_ experiments, cells were incubated with 2 mM MitoPY1 (without replacing the cells’ medium) for 1 h, at the same experimental conditions. Then, cells were washed 1x with warm PBS and placed in Krebs solution, and the fluorescence signal was read for 30 min (λEx = 503 nm and λEm = 540 nm), at 37°C, using a plate reader (Fluorimeter SpectraMax Gemini EM, Molecular Devices, San Jose, CA, USA).

The protein content of each well was determined by aspirating the assay buffer, followed by addition of 20 µL RIPA buffer. The multi-well plate was placed on a plate shaker at low speed, for 10 min, and then on ice during 20 min. Finally, 200 µL BCA assay mix were added. After a 30 min incubation period, absorbance was measured at 562 nm with the microplate reader spectraMax iD3.

### 2.8. Colorimetric determination of thiobarbituric acid reactive substances (TBARS) levels

The thiobarbituric acid (TBA) assay was used to measure TBARS levels at basal conditions, as previously described [41]. Briefly, 250 µL of reaction medium (175 mM KCl, 10 mM Tris-HCl, pH 7.4) was added to 150 µg of protein and incubated at 37°C. To start the reaction, 1 mM ADP/0.1 mM Fe^2+^ were added to each sample and after 15 min of incubation 250 µL of 40% (*w/v*) trichloroacetic acid (TCA) was added to stop the reaction. Then, 1 mL of 0.67% (*w/v*) TBA was added, vortex-mixed, and boiled at 100°C for 15 min. Samples were centrifuged for 10 min at 18,000 x g and the absorbance of the supernatants was measured at 530 nm with a Spectrophotometer Spectramax plus 384. The amount of TBARS was calculated using a molar coefficient of 1.56 x 10^5^ M/cm and expressed as nmol TBARS/mg protein.

### 2.9. Fluorometric analysis of mitochondrial membrane potential

To evaluate the mitochondrial membrane potential (Δψm) at basal conditions, the fluorescent probe tetramethylrhodamine ethyl ester perchlorate (TMRE) was used. Cells were cultured in 96-well plate (6,250 cells per well), in triplicate, and incubated in a humidified 5% CO_2_-95% air atmosphere at 37°C. After 72 h of incubation, the culture medium was changed by fresh medium containing 4 µM TMRE (Sigma-Aldrich, 87917) and incubated during 30 min at 37°C. Then, cells were washed 1x with warm PBS and the fluorescence signal (λEx = 544 nm; λEm = 590 nm) was read at 37°C using a microplate reader spectraMax iD3 (Molecular Devices, San Jose, CA, USA).

To determine the protein concentration of each well, the assay buffer was removed and 10 µL of RIPA buffer was added to each sample. The multi-well plate was placed on a plate shaker at low speed for 10 min and then on ice for 20 min. For protein quantification, 100 µL BCA assay mix were added, and, after a 30 min of incubation, the absorbance was read at 562 nm with a microplate reader spectraMax iD3.

### 2.10. Lipidomic analysis

#### 2.10.1. Subcellular fractionation

For the lipidomic analysis, microsomes and MAM were obtained as previously described [25]. Briefly, to remove cell debris and nuclei and obtain the total fraction (TF), cells were homogenized in isolation buffer (250 mM sucrose and 10 mM HEPES, supplemented with a 1% cocktail of proteases inhibitors) and centrifuged at 600 x g for 5 min, at 4°C. After, the TF was centrifuged at 4°C, for 10 min, at 10,300 x g, to obtain the supernatant containing the ER/microsomal fraction and the pellet containing the MAM fraction. To pellet the ER/microsomal fraction, the supernatant was centrifuged for 60 min, at 100,000 x g at 4° C, in a Beckman ultracentrifuge (Indianapolis, Indiana, USA, model L-100 XP, 90 Ti rotor). The MAM fraction was centrifuged at 95,000 x g during 65 min, at 4°C, in a Beckman ultracentrifuge (model L-100 XP, SW41 rotor) after layering on top of a 30% (*v/v*) Percoll gradient. To pellet the MAM fraction, the upper band (containing MAM fraction) was diluted with PBS containing 1 mM PMSF and centrifuged at 6,300 x g, for 10 min, at 4°C. The supernatant was centrifuged during 30 min, at 100,000 x g, at 4°C, in a Beckman ultracentrifuge (model L-100 XP, 90 Ti rotor). The purity of each subcellular fraction was validated by Western blot (WB), as previously described [25].

#### 2.10.2. Lipids extraction

The total lipid content of each subcellular fraction was extracted according to the Bligh and Dyer method, as previously described [42]. The subcellular fractions were resuspended in 1 mL of water, transferred to glass tubes, followed by the addition of 3.75 mL dichloromethane/methanol 1:2 (*v/v*) and incubation on ice for 30 min, with occasional vortex. After, 1.25 mL of dichloromethane and 1.25 mL Milli-Q H_2_O was added and tubes were centrifuged at 2,000 rpm (Mixtasel centrifuge, Selecta) at RT, during 10 min. The organic phase was collected into a new glass tube and the aqueous phase was re-extracted 3x with 1.88 mL dichloromethane. Combined organic phases of each sample were dried under a nitrogen stream and stored at -20°C.

#### 2.10.3. Quantification of phospholipid content by the phosphorus assay

The total amount of phospholipids was quantified as previously described [42,43]. Briefly, the phosphorus present in 10 μL of dried total lipid extracts, previously dissolved in 300 μL dichloromethane, were incubated with 70% (*v/v*) 125 μL perchloric acid in a heating block (Stuart, UK) during 1 h, at 180°C. Samples were vortex mixed after each addition of 825 μL of Milli-Q water, 125 μL of 2.5% (v/v) (NH_2_)_6_MoO_4_·4H_2_O, and 125 μL of 10% (*v/v*) ascorbic acid and incubated at 100°C in a water bath for 10 min. The absorbance of samples and standards of 0.1-2 μg (NaH_2_PO_4_⋅2H_2_O, 100 μg of phosphorus mL^−1^), which were prepared in the same way as samples, except the heating block step, were read at 797 nm using a Multiskan GO1.00.38 Microplate Spectrophotometer (Thermo Scientific, Hudson, NH, USA) controlled by SkanIT software, version 3.2 (Thermo Scientific) [42].

#### 2.10.4. Lipid analysis using C18 liquid chromatography – mass spectrometry (LC-MS)

Reverse phase liquid chromatography was performed in an Ultimate 3000 Dionex (Thermo Fisher Scientific, Bremen, Germany) coupled to the Q-Exactive® hybrid quadrupole Orbitrap mass spectrometer (Thermo Fisher, Scientific, Bremen, Germany), as previously described [44–46]. An amount of 10 µg of phospholipids of each sample (in 10 µL of dichloromethane) were mixed with 82 µL of 50% isopropanol/ 50% methanol and 8 µL of phospholipid standards mixture (dMPC - 0.04 µg, SM d18:1/17:0 - 0.04 µg, dMPE - 0.04 µg, LPC - 0.04 µg, dPPI - 0.08 µg, CL(14:0)4 - 0.16 µg; dMPG - 0.024 µg, Cer 17:0/d18:1 - 0.08 μg, dMPS - 0.08 µg; dMPA-0.16 μg). A volume of 5 µL was loaded into the C18 column (Sigma-Aldrich®, 2.1 x 100 mm, 2.7 µm) at 50°C and at a flow-rate of 260 µL min^−1^. The gradient applied was: 32% B at 0 min, 45% B at 1.5 min, 52% B at 4 min, 58% B at 5 min, 66% B at 8 min, 70% B at 11 min, 85% B at 14 min, 97% B at 18 min, 97% B at 25 min, 32% B at 25.01 min and 32% B at 33 min. Mobile phases A and B were composed of water/acetonitrile (40/60%) and isopropanol/acetonitrile (90/10%), respectively, with 10 mM ammonium formate and 0.1% formic acid. The mass spectrometer operated using positive/negative switching toggles between positive (ESI+, 3.0 kV) and negative (ESI-, 2.7 kV) ion modes, with the following parameters: capillary temperature of 320°C; sheath gas flow of 35 U; high resolution of 70,000, automatic gain control (AGC) of 3 x 10^6^, maximum inject time (IT) of 100 ms, *m/z* range of 200-1600, and 2 microscans for full scan mode; resolution of 17,500, AGC target of 1x10^5^, maximum IT of 100 ms, and 1 microscan for tandem MS. The data-dependent method included up to 10 MS/MS spectra per full-scan mass spectrum. Cycles were repeated with a dynamic exclusion of 30 s and an intensity threshold of 8 x 10^4^. Normalized collision energyTM (CE) used for MS/MS acquisition ranged between 20, 24 and 28 eV in ESI- and 25 and 30 eV in ESI+. Xcalibur data system (V3.3, Thermo Fisher Scientific, Bremen, Germany) was used for data acquisition.

#### 2.10.5. Lipidomic data analysis

Lipid species were identified based on mass accuracy observed in C18-RP-HPLC-ESI-MS and MS/MS spectra interpretation. Raw data import, feature detection and peak area integration were performed using Lipostar software (Molecular Discovery Ltd., version 2.1.4 × 64) [47]. Raw files were imported applying an automatic MS signal threshold of 100. Feature detection was carried out using *m*/*z* tolerance of ±0.02 Da; signal filtering threshold of 10000; MS/MS filtering threshold and chromatogram filtering threshold of 0.97. Lipid assignment and identification were made against a database generated from the LIPID MAPS structure database (version March 2024). This database was further fragmented using the DM Manager Module within Lipostar software, adhering to its fragmentation rules. The raw files were directly imported and aligned, following the settings specified by [48]. Automatic peak picking was performed with SDA smoothing level set to high and a minimum S/N ratio of 3 and an automated approval was executed to retain structures rated between 2 to 4 stars in terms of quality. For the lipid identification the parameters of 5 ppm precursor ion mass tolerance and 20 ppm product ion mass tolerance were followed. The identification of each lipid species was based on the *m/z* value of the ions identified in the LC-MS data, exact mass measurements (mass accuracy ≤ 5 ppm), retention time, and MS/MS data analysis. Relative quantification was performed by exporting the Integrated peak area values from the lipid species in the comma separated values (.csv) format. The extracted-ion chromatogram (XIC) was obtained through the sum of the areas of all identified lipid species of DAG or TAG, and the data normalization was performed by dividing the XIC peak by the total content of phospholipids in the lipid extract.

### 2.11. Statistical analysis

Statistical analysis was performed using the GraphPad Prisma 8.0.2 software (San Diego, California, USA) and presented as mean ± standard error of the mean (SEM). All experiments were independent assays, with each *n* representing a biological replicate, calculated as the average of 3 technical replicates or, in the case of microscopy, the average of 5 fields of view. Data were tested for Gaussian distribution using the Shapiro-Wilk test. Differences between the two groups were analyzed using unpaired *t*-test and two-way ANOVA with Sidak post hoc correction for grouped analysis. In cases of non-normality distribution, data were analyzed by the Mann-Whitney U-test to compare the two groups. Statistical significance was considered at *p* ≤ 0.05.

## 3. Results

### 3.1. LD changes are associated with altered FFA and cholesteryl ester concentrations in APPswe cells

We have previously reported a decrease in the close ER-mitochondria contacts associated with mitochondrial dysfunction [25] and an altered lipidome in APPswe cells [42]. Knowing that MAM is enriched in proteins involved in LD metabolism [2], we characterized LD by confocal microscopy using the LipidTox Red probe. A decrease in the number (p<0.001) (Figure 1a and b), and an increase in the area and volume (p<0.001) (Figure 1c and d) of LD were observed in APPswe cells. Since these observations suggest an impairment in lipid storage in APPswe cells, we measured the concentration of FFA, total cholesterol, and CE using a commercial kit. We observed an increase in FFA (p<0.05) and a decrease in total cholesterol (p<0.001) and CE (p<0.05) concentration in APPswe cells (Figure 1e-g). Interestingly, WB analyses revealed that the protein levels of acetyl-CoA carboxylase 1 (ACC1), which is involved in *de novo* FA synthesis [49], is increased (p<0.001) while those of ACAT1/SOAT1, which catalyzes the esterification of cholesterol into CE [50], is decreased (p<0.01) in APPswe cells (Figure 1h-j). Altogether, these findings suggest an altered metabolism of LD in APPswe cells. Since we observed decreased LD and enhanced FFA, we hypothesized that an increase of LD degradation and/or an impairment in LD biosynthesis may occur in APPswe cells.

**Figure 1.**
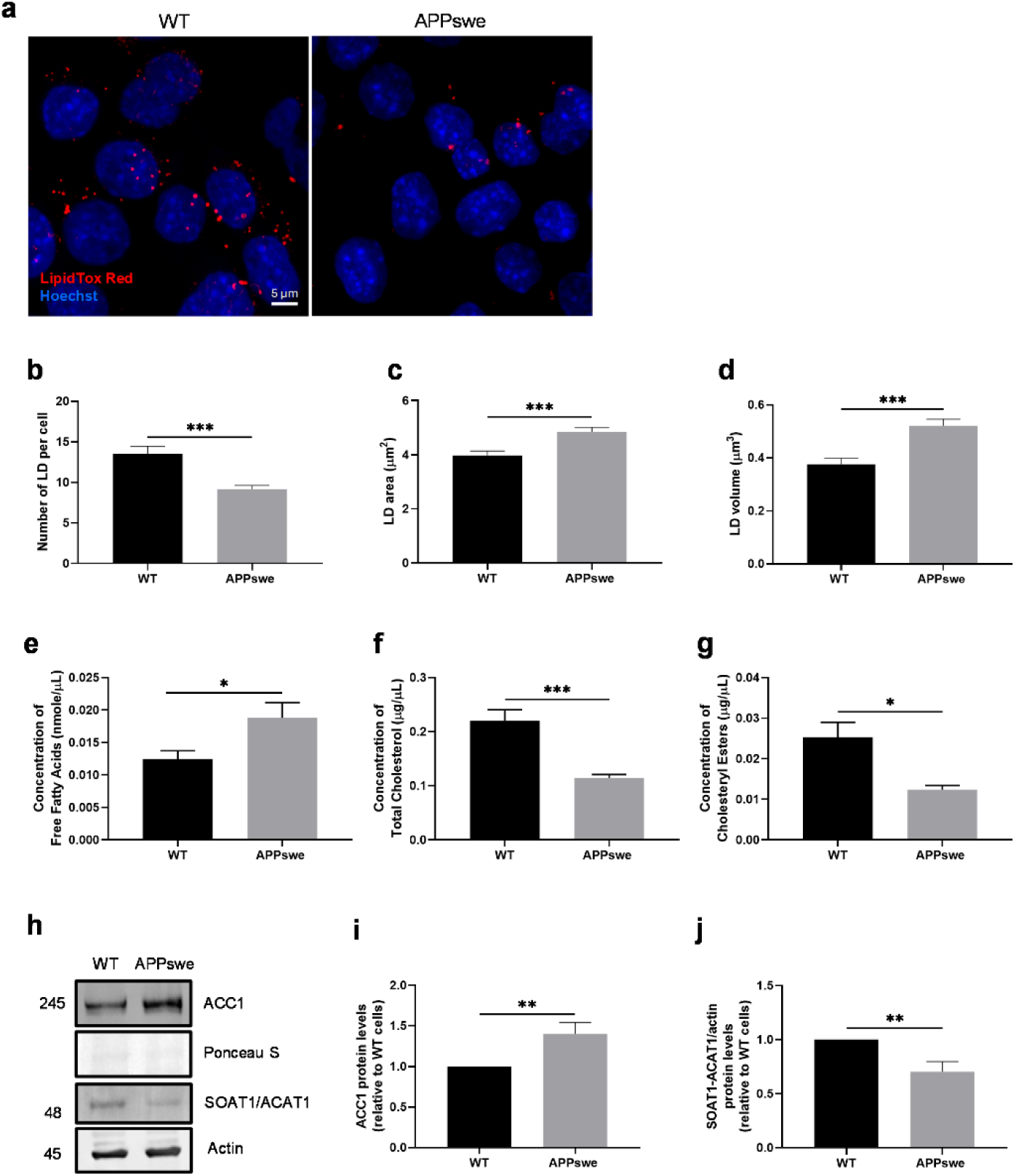
Altered lipid droplet (LD) characteristics and lipids levels in APPswe cells. (a) Representative image of LipidTox Red (LD, red) and Hoechst (nucleus, blue) staining. Scale bar represents 5 µm. (b-d) Quantification of LD number normalized by cell number, area, and volume (n=4) (data from cells treated with DMSO during 48h). (e) Quantification of free fatty acid (FFA) concentration (n=6). (f) Quantification of total cholesterol and (g) cholesteryl ester (CE) concentration (n=5-6). (h-j) Representative Western blots (WB) and quantification of the acetyl-CoA carboxylase 1 (ACC1) and acyl-CoA: cholesterol acyltransferase 1/sterol O-acyltransferase 1 (ACAT1/SOAT1) (n=6-8). All data is presented as mean ± SEM; *p*-values were obtained using the unpaired *t*-test and non-parametric independent Mann-Whitney U test. * *p* ≤ 0.05, ** *p* ≤ 0.01, and *** *p* ≤ 0.001 were considered statistically significant.

### 3.2. Macroautophagy is altered in APPswe cells but does not play a significant role in LD alterations

FA released from LD, upon autophagy-mediated degradation, can be used for ATP production by mitochondrial β-oxidation, which produces substrates for the tricarboxylic acid (TCA) cycle [7,15,51] (Figure 2a). In this context, we analyzed the interaction between LD and the lysosomal marker Lamp1. An increase of LD-Lamp1 co-localization (p<0.001) (Figure 2b and c), and in the number and volume of lysosomes (p<0.01) were found in APPswe cells, when compared with WT cells (Figure 2d and e), suggesting an increased LD degradation by macroautophagy. Next, we evaluated the co-localization of LD and the autophagosome marker LC3, because LC3-positive autophagic membranes surround LD and promote the entry of LD into autophagosomes (Figure 2a) [15]. In APPswe cells, we found an increased LC3-LD co-localization when compared to WT cells (Figure 2b and f). When we treated the cells with CQ, which inhibits the autophagic flux causing an accumulation of LC3-II [52] by decreasing the autophagosome-lysosome fusion [53], we observed, by WB, an accumulation of LC3-II (p<0.001) in both WT and APPswe cells (Figure 2g). However, a modest decrease of the autophagic flux is observed in APPswe cells when compared to WT cells (Figure 2h), suggesting an impairment of macroautophagy in APPswe cells. Interestingly, no significant alterations in LD number in both WT and APPswe cells were observed upon CQ treatment, when compared with non-treated cells. Nevertheless, a reduction of LD in APPswe cells occurred when compared with WT cells (p<0.0001) (Figure 2i and j). To understand the role of macroautophagy in LD degradation under starvation, which stimulates autophagy, we treated the cells with OA, to promote LD formation, and with CQ, under starvation, to induce LD degradation. Under these experimental conditions, we observed an increase of LD number in WT (p<0.0001) and APPswe (p<0.05) cells in the presence of CQ (Figure 2k and l); however, the increase was more pronounced in WT cells than in APPswe cells, which suggests that macroautophagy can degrade LD under starvation in both cell lines but cannot be responsible for the reduced number of LD in APPswe cells. Thus, macroautophagy is compromised in APPswe cells, however, this quality control process does not seem to play a major role in the LD alterations found in APPswe cells.

**Figure 2.**
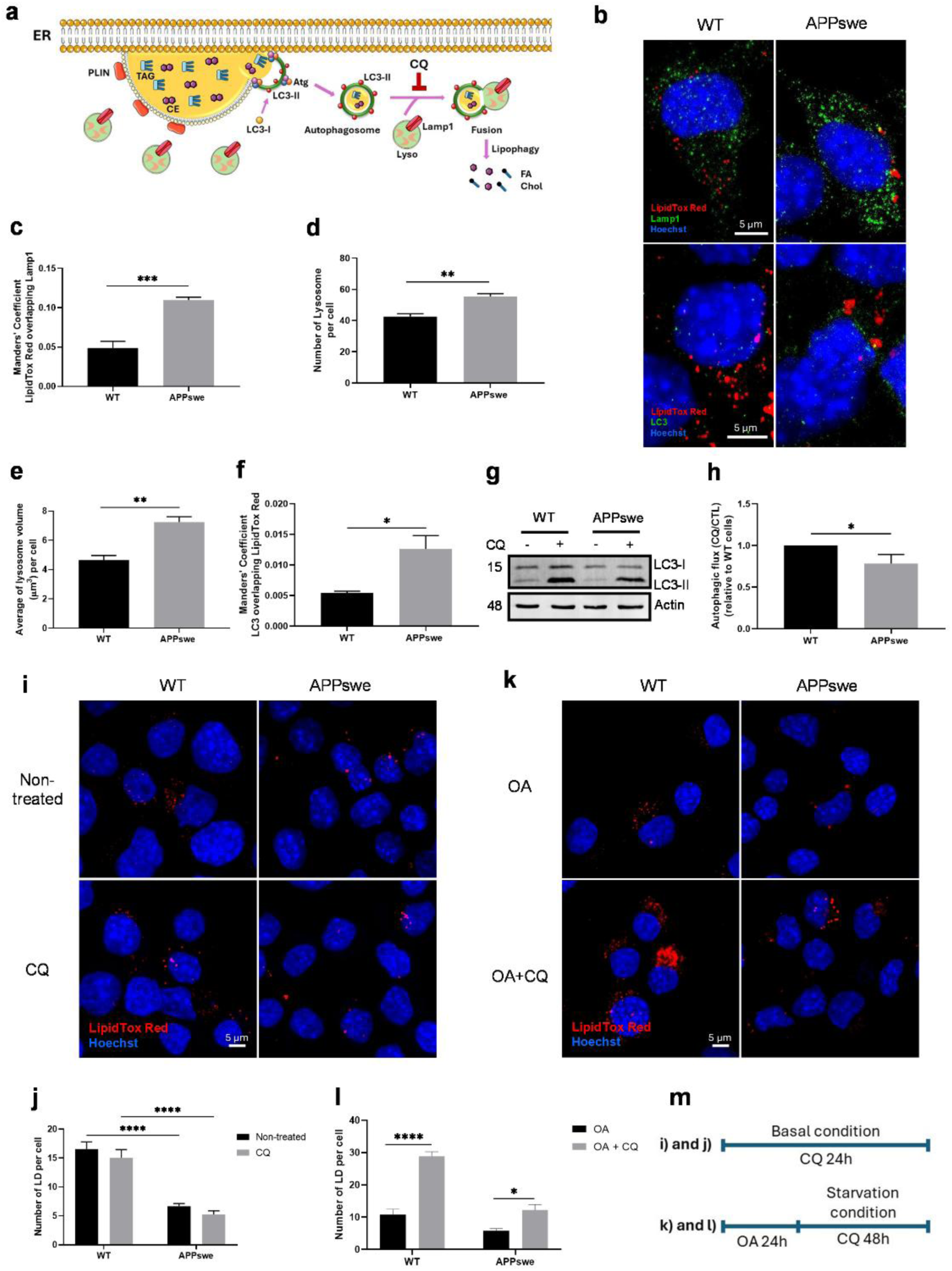
Macroautophagy effects on lipid droplets (LD) in APPswe cells. (a) The cartoon shows macroautophagy-mediated LD degradation and the blocking effect mediated by chloroquine (CQ). (b) Representative image of LipidTox Red (LD, red), lysosome-associated membrane protein type 1 (Lamp1) or microtubule-associated protein light chain 3 (LC3) (green), and Hoechst (nucleus, blue). Scale bar represents 5 µm. (c) Quantification of Manders’ coefficient of LD overlapping Lamp1 (n=4). (d) Quantification of lysosome number normalized by cell number (n=3) and (e) mean of lysosome volume normalized by cell number (n=3). (f) Quantification of Manders’ coefficient of LC3 overlapping LD (n=3). (g) Representative WB of LC3-I and LC3-II proteins and (h) quantification of autophagic flux (ratio between cells treated for 24 h with 20 µM CQ and non-treated cells) (n=7-8). (i) Representative image of LipidTox Red (LD, red) and Hoechst (nucleus, blue) staining in cells treated for 24 h with 20 µM CQ and (j) quantification of LD number (n=4). Scale bar represents 5 µm. (k) Representative image of LipidTox Red (LD, red) and Hoechst (nucleus, blue) staining in cells treated for 24 h with 25 µM oleic acid (OA) plus 10 µM CQ during 48 h under starvation and (l) quantification of LD number normalized by cell number (n=4). (m) The cartoon shows the treatment conditions for i-k images. All data presented as mean ± SEM; *p*-values were obtained using unpaired *t*-test. * *p* ≤ 0.05, ** *p* ≤ 0.01, *** *p* ≤ 0.001, and **** *p* ≤ 0.0001 were considered statistically significant.

### 3.3. CMA, but not the ubiquitin-proteosome system (UPS), is involved in LD reduction in APPswe cells

The interaction between LD and lysosomes allows the degradation of PLIN2 and PLIN3, proteins that surround LD membranes, via CMA. In this process, hsc70 recognizes the pentapeptide motif of PLIN2 and PLIN3, delivers these proteins to the lysosome surface facilitating their translocation to the lysosomal lumen through the receptor Lamp2A [2,7]. To study the role of CMA in LD degradation, we analyzed the number of LD and PLIN2-Lamp2A interaction under different experimental conditions, namely under basal conditions, starvation, OA treatment, and OA treatment followed by starvation.

A significant reduction in LD number in APPswe cells was found in all experimental conditions when compared with WT cells (Figure 3a and b). Of note, we observed a decrease of LD number (p<0.0001) and an increase of PLIN2-Lamp2A (p<0.05) interaction in APPswe cells under basal conditions when compared to WT cells (Figure 3c), suggesting that enhanced CMA activity may contribute to the reduction in LD number observed in APPswe cells.

**Figure 3.**
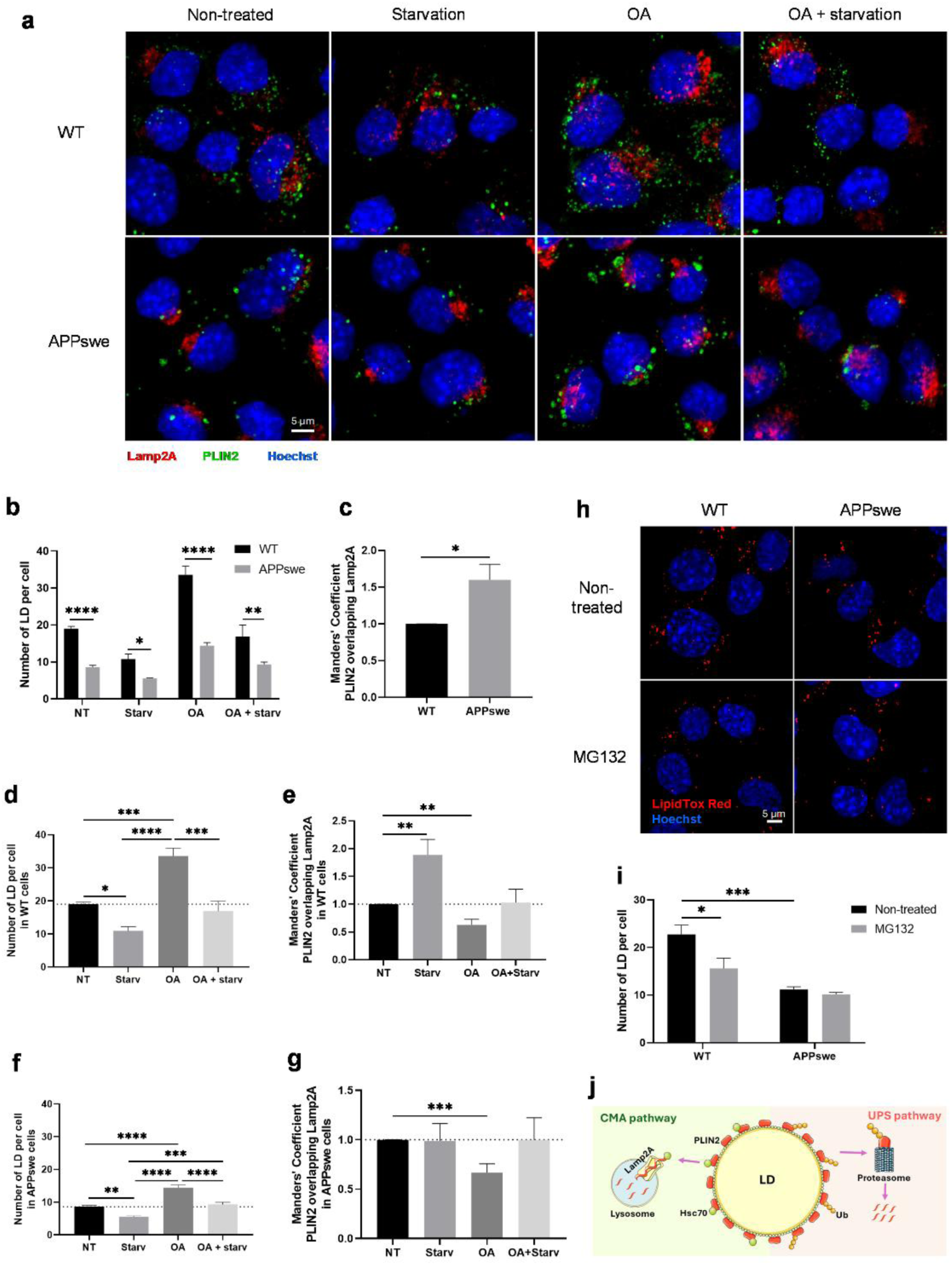
Involvement of chaperone-mediated autophagy (CMA) in lipid droplets (LD) degradation in the APPswe cells. (a) Representative image of lysosome-associated membrane protein type 2A (Lamp2A) (lysosome, red), perilipin 2 (PLIN2) (green), and Hoechst (nucleus, blue) staining in non-treated cells, upon 24 h of starvation, treated with 25 µM oleic acid (OA) for 24 h, and treated with 25 µM OA for 24 h plus 24 h of starvation. Scale bar represents 5 µm. (b) Quantification of LD in WT and APPswe cells normalized by cell number (n=6). (c) Quantification of Manders’ coefficient of PLIN2 overlapping Lamp2A in non-treated (NT) cells relative to WT cells (n=5-6). (d) Quantification of LD number normalized by cell number and (e) quantification of Manders’ coefficient of PLIN2 overlapping Lamp2A relative to NT WT cells (n=6). (f) Quantification of LD number normalized by cell number and (g) Manders’ coefficient of PLIN2 overlapping Lamp2A, relative to NT APPswe cells (n=6). (h) Representative image of LipidTox Red (LD, red) and Hoechst (nucleus, blue) staining in cells exposed to 50 nM MG132 for 24 h and (i) quantification of LD number normalized by cell number (n=4). Scale bar represents 5 µm. (j) The cartoon shows PLIN2 degradation via CMA and ubiquitin proteosome system (UPS) pathways. All data presented as mean ± SEM; *p*-values were obtained using unpaired *t*-test and two-way ANOVA with Sidak’s multiple comparisons test. * *p* ≤ 0.05, ** *p* ≤ 0.01, *** *p* ≤ 0.001, and **** *p* ≤ 0.0001 were considered significant.

In WT cells, the number of LD decreases under starvation (p<0.05), increases after OA treatment (p<0.001), and remains unaltered after OA treatment followed by starvation, when compared with the basal conditions. As expected, we also observed a decrease in LD number in cells under starvation after OA treatment, when compared with cells treated with OA only (p<0.001) (Figure 3d). To study the role of CMA in LD degradation, we analyzed the PLIN2-Lamp2A interaction, and we found an increased co-localization of these proteins in WT cells under starvation (p<0.01) and a decreased co-localization in cells treated with OA (p<0.01), when compared to basal conditions (Figure 3e). These results show that in WT cells the reduction in LD number, under starving conditions, is associated with an increased PLIN2-Lamp2A co-localization, suggesting that the degradation of LD occurs through CMA in these cells.

A similar profile was found in APPswe cells exposed to the above-mentioned experimental conditions. In APPswe cells, we found a decrease in LD number under starvation (p<0.01), an increase of LD number after OA treatment (p<0.0001), and a reduction in LD number to baseline values after OA treatment followed by starvation, when compared with APPswe cells under basal conditions. As observed in WT cells, the APPswe cells revealed a decrease in LD number under OA treatment followed by starvation, when compared with APPswe cells treated with OA only (p<0.0001) (Figure 3f). However, in APPswe cells we only observed a decrease in PLIN2-Lamp2A co-localization under OA treatment (p<0.001), when compared to APPswe cells exposed to basal conditions (Figure 3g). Altogether, these results suggest that CMA can be activated in APPswe cells and can be involved in PLIN2 protein degradation under basal conditions, but not under starvation. Furthermore, our results also suggest that the CMA does not play a major role in LD degradation under OA treatment in both cell lines (Figure 3d-g).

UPS is one of the major protein degradation pathways [54], which can be involved in PLIN2 degradation [55], and UPS inhibition can promote the degradation of LD by the autophagy-lysosome pathway [54]. We analyzed the content of LD in the presence of the proteasome inhibitor MG132, to clarify if this pathway is involved in LD degradation. We found a decrease in LD number in WT cells treated with MG132 (p<0.05); however, no alterations were observed in APPswe cells in the presence of this inhibitor (Figure 3h and i), suggesting that the UPS does not play an important role in LD degradation in APPswe cells.

Altogether, our results suggest that CMA may be involved in the degradation of PLIN2 and promote TAG lipolysis by increasing FFA levels in APPswe cells.

### 3.4. Decreased seipin levels correlate with LD and lipid dyshomeostasis and a pro-oxidative environment in APPswe cells

Our next step was to investigate the ability of cells to synthesize LD. Cells were treated with OA, which promotes LD biogenesis. We observed that OA treatment increases LD number in WT (p<0.0001) and APPswe cells (p<0.01). However, in APPswe cells the increase in the number of LD was less pronounced (p<0.0001), and their size was more heterogeneous, when compared with WT cells (Figure 4a and b), suggesting an impaired LD biosynthesis in APPswe cells.

**Figure 4.**
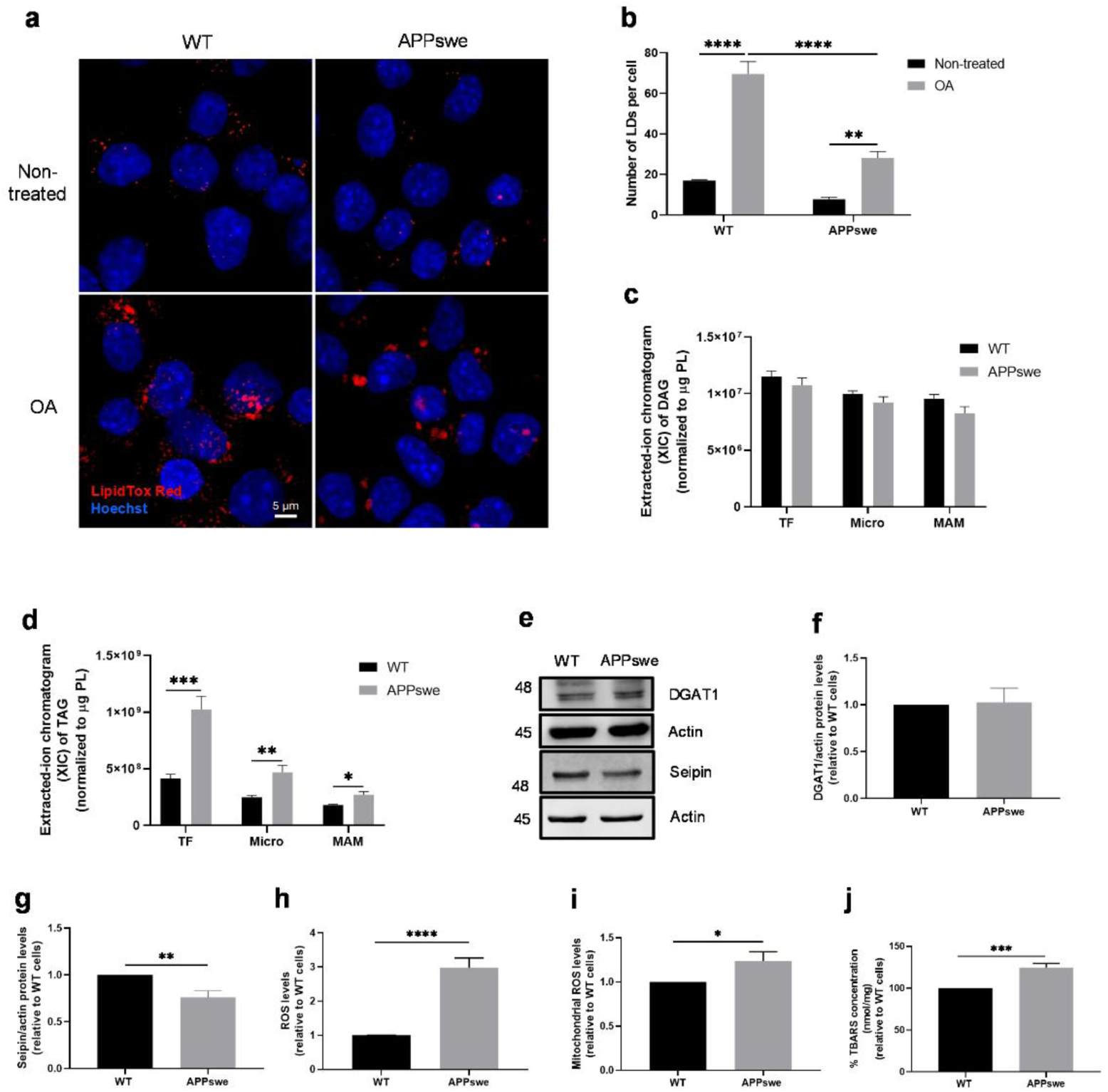
Lipid droplet (LD) biogenesis, diacylglycerol (DAG) and triacylglycerol (TAG) amount and oxidative status in APPswe cell line compared with WT cell line. (a) Representative image of LipidTox Red (LD, red) and Hoechst (nucleus, blue) staining in cells treated with 25 µM oleic acid (OA) for 24 h and (b) quantification of LD normalized by cell number (n=4). Scale bar represents 5 µm. (c-d) Relative amount of DAG and TAG using extracted-ion chromatogram (XIC) in different subcellular fractions: total fraction (TF), microsomes (Micro), and mitochondria-associated membranes (MAM) (n=6). (e) Representative WB and quantification of (f) DAG acyltransferase 1 (DGAT1) and (g) seipin levels (n=7-8). (h) Quantification of reactive oxygen species (ROS) using the dichlorodihydrofluorescein diacetate (DCFH2-DA) probe (n=6). (i) Quantification of mitochondrial ROS using the MitoPY1 probe (n=8). (j) Quantification of thiobarbituric acid reactive substances (TBARS) using the TBA assay (n=8). All data presented as mean ± SEM; *p*-values were obtained using unpaired *t*-test. * *p* ≤ 0.05, ** *p* ≤ 0.01, *** *p* ≤ 0.001, and **** *p* ≤ 0.0001 were considered statistically significant.

LD biogenesis begins with the synthesis of neutral lipids, most commonly TAG and sterol esters that result from the esterification of an activated fatty acid to DAG or a sterol, respectively [7]. Therefore, we analyzed by reversed phase liquid chromatography coupled to mass spectrometry the amount of DAG and TAG in different subcellular fractions: total fraction, microsomes, and MAM. Our analyses did not show any significant difference in the relative amount of DAG in all fractions between the two cell lines (Figure 4c), but a significant increase in the relative amount of TAG in total fraction (p<0.001), microsomes (p<0.01), and MAM (p<0.05) was observed in APPswe cells, when compared with WT cells (Figure 4d). Then, we measured by WB the protein levels of DGAT1, that promotes the esterification of DAG in TAG, but no significant differences were found between the two cell lines (Figure 4e and f). However, DGAT1 activity can be affected without any change in protein levels. The increase of TAG in APPswe (Figure 4d) together with the decrease of LD number, despite the increase of LD area and volume (Figure 1c and d), suggest an impairment in LD biogenesis in APPswe cells.

Seipin is an ER transmembrane protein that is enriched at close ER-mitochondria contacts (*i.e.*, at MAM) and is involved in LD biogenesis [10,11]. It is also described that at later stages of LD biogenesis, each LD is associated with, at least, one seipin punctum and this protein also supports the ER-LD contact formation and promotes the incorporation of TAG into LD [9,11,56]. Since we observed an alteration in LD number and size, concomitantly with an increase of TAG in APPswe cells, we analyzed, by WB, the protein levels of seipin. A significant decrease in the levels of this protein was found in APPswe cells, when compared with WT cells (p<0.01) (Figure 4e and g). In summary, these results suggest that seipin depletion affects LD biogenesis, resulting in fewer but larger LD in APPswe cells, being LD expansion associated with TAG accumulation in these cells.

Under physiological conditions, cells produce low to moderate levels of ROS that play an important role in cell signaling and homeostasis. On the other hand, the overproduction of ROS can induce deleterious alterations in cellular components, including lipids [57,58]. We measured the cytosolic and mitochondrial ROS levels using the DCFH2-DA and MitoPY1 fluorescence probes, respectively, and we found an increase in both cytosolic (p<0.0001) and mitochondrial (p<0.05) ROS levels in APPswe cells, compared to WT cells (Figure 4h and i). Since our results demonstrate an accumulation of FFA, which are susceptible to ROS damage, we evaluated the extent of lipid peroxidation by measuring the levels of TBARS, a product of lipid peroxidation [41]. We observed an increase of ∼25% in TBARS levels in APPswe cells, when compared with WT cells (p<0.001) (Figure 4j). Taking together, our findings suggest that lipid oxidation can result from increased ROS levels associated with an impaired storage of lipids into LD leading to FFA accumulation in APPswe cells.

### 3.5. DGAT1 inhibition recovers the ER-mitochondria contacts at MAM in APPswe cells

We have observed an increase of TAG levels (Figure 4d), which are normally stored in LD to prevent lipotoxicity and to provide energy substrates to the mitochondria. Since DGAT1 is an enzyme involved in TAG synthesis and essential to remodel and finish the LD biogenesis process [59], we treated both WT and APPswe cells with A922500, a DGAT1 inhibitor that prevents TAG synthesis (Figure 5a). In the absence of the inhibitor A922500, we found a decrease in the number of LD and an increase in area and volume of these organelles (p<0.001) in APPswe cells, when compared with WT cells. As expected, WT and APPswe cells treated with the inhibitor A922500 showed a lower number of LD (p<0.0001 and p<0.001, respectively) (Figure 5b and c). No significant alterations in LD size were found in WT cells exposed to A922500, while APPswe cells showed a decrease in LD area and volume under similar conditions (p<0.01) (Figure 5d and e). These results suggest a recovery of LD size in APPswe cells upon DGAT1 inhibition.

**Figure 5.**
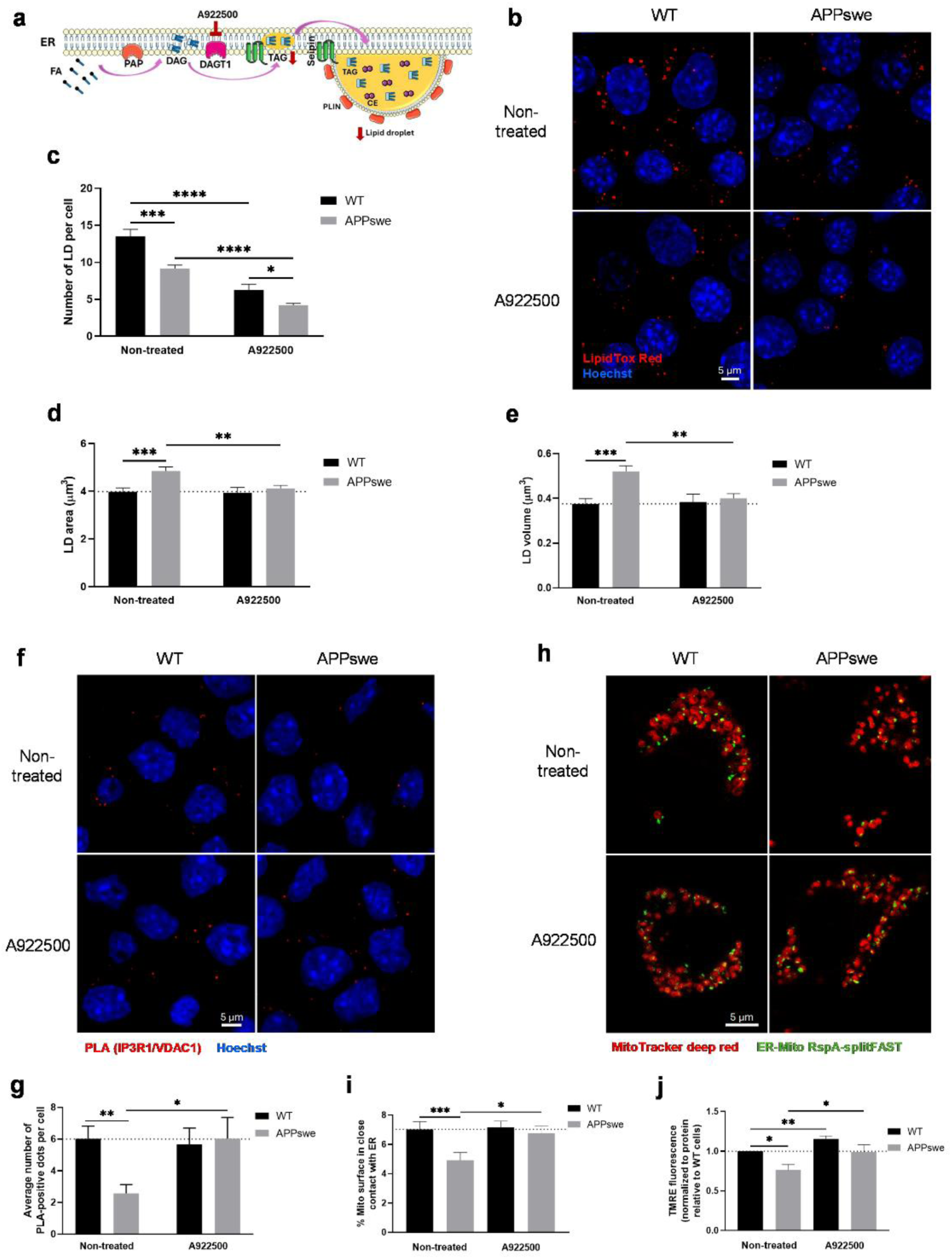
Inhibition of diacylglycerol acyltransferase 1 (DGAT1) increases close ER-mitochondria contacts and improves mitochondrial function in the APPswe cells. (a) The cartoon shows the lipid droplet (LD) formation and A922500 targeting. (b) Representative image of LipidTox Red (LD, red) and Hoechst (nucleus, blue) staining in cells treated with 20 µM A922500 for 48 h and (c-e) analysis of LD number normalized by cell number, area, and volume (n=4). The data presented in non-treated condition was already presented in Figure 1b-d. Scale bar represents 5 µm. (f) Representative image of proximity-ligation assay (PLA) using anti-IP3R1 and -VDAC1 antibodies (PLA, red) and Hoechst (nucleus, blue) staining in cells treated with 20 µM A922500 for 48 h and (g) quantification of the average number of PLA-positive dots normalized to number of cells (n=6). Scale bar represents 5 µm. (h) Representative image of MitoTracker deep red (mitochondria, red) and ER-mitochondria contact RspA-splitFAST (ER-mitochondria contacts, green) staining in cells treated with 20 µM A922500 for 48 h and (i) quantification of the percentage of mitochondria surface in close contact with the ER (n>104 cells from 3 biological replicates). Scale bar represents 5 µm. (j) Quantification of mitochondrial membrane potential (Δψm) using the tetramethylrhodamine ethyl ester perchlorate (TMRE) fluorescence probe (n=8). All data presented as mean ± SEM; *p*-values were obtain using unpaired *t*-test, non-parametric independent Mann-Whitney U test, and two-way ANOVA with Sidak’s multiple comparisons test. * *p* ≤ 0.05, ** *p* ≤ 0.01, *** *p* ≤ 0.001, and **** *p* ≤ 0.0001 were considered statistically significant.

Lipid alterations can affect protein-protein interactions [60] and accumulation of lipids can disrupt the ER endomembrane homeostasis [61,62]. Previously, we observed a decrease in MAM and an impairment of mitochondrial function, including mitochondrial membrane depolarization and altered respiration in APPswe cells, when compared with WT cells [25]. To investigate if the decrease of TAG synthesis affects ER-mitochondria contacts, we used the PLA assay, and the new ER-mitochondria contact RspA-splitFAST probe [39] to analyze the physical interaction between both organelles in WT and APPswe cells exposed to A922500. Under basal conditions, we observed a decrease in the close ER-mitochondria contacts in APPswe cells when compared with WT cells (p<0.05 in PLA assay and p<0.001 with the ER-mitochondria contact RspA-splitFAST probe), which is in accordance with our previous findings [25]. However, we found an increase in these contacts in APPswe cells treated with A922500 (p<0.05), a situation that was not observed in WT cells (Figure 5f-i). Using the TMRE probe, we observed a loss of Δψm in APPswe cells when compared with WT cells (p<0.05), under basal conditions. Interestingly, this mitochondrial parameter was improved in WT (p<0.001) and APPswe (p<0.05) cells treated with A922500. Of note, APPswe cells exposed to A922500 presented a Δψm value like that found in WT cells under basal conditions (Figure 5j). In conclusion, these results suggest that DGAT1 inhibition and subsequent decrease of TAG synthesis restores the contacts between the ER and mitochondria and mitochondrial polarization in APPswe cells.

## 4. Discussion

Alterations in lipid storage have been associated with pathological conditions, such as obesity, inflammation, cancer, and neurodegenerative diseases [15]. Lipotoxicity, mitochondrial dysfunction, and increased oxidative stress have been linked with insufficient lipid storage, while hypoxia, ER stress, immune cells infiltration, and increased secretion of proinflammatory cytokines have been related to excessive lipid accumulation [28,63]. Changes in different lipid classes, such as cholesterol, sphingolipids, phospholipids, and glycerolipids were reported in AD preclinical models and in samples from patients, including plasma, brain, and cerebrospinal fluid [64,65].

We have previously shown alterations in close ER-mitochondria contacts [25] and in lipid homeostasis in different subcellular fractions (ER, mitochondria, and MAM) in the *in vitro* neuroblastoma model of AD used in the present study [42]. Here, we observed a decrease of LD number and an increase of FFA concentration. Although FA are an important energy source for cells, increased levels can be toxic. Therefore, the storage of FA into LD prevents lipotoxicity by working as a cell buffering system [7]. An increase in LD number in fibroblasts from FAD and SAD patients, PSEN1-mutant MEFs [28], 3xTg-AD mouse model, and AD human brain tissue [66] has been reported. Increased LD number has been associated to elevated CE levels and ACAT1/SOAT1 activity in PS-mutant MEFs as well as to elevated CE synthesis in PS-mutant APPswe cells [28]. In the present study, we found a decrease in LD number associated with lower total cholesterol and CE levels, and ACAT1/SOAT1 protein levels. The inhibition of ACAT1/SOAT1 activity caused a lower concentration of CE in both liver and brain of transgenic mice expressing the human APP_751_ containing the London (V717I) and Swedish (K670M/N671L) mutations [67] and an accumulation of cholesterol at MAM [50]. However, the decrease in LD number and the increase of FFA observed in the present study with APPswe cells could be associated with the catabolism of TAG stored in LD to generate FFA through the action of cytosolic [68] and lysosomal lipases by the sequestration of complete or small portions of LD by macrolipophagy [69,70]. We also found an increase in LD-lysosome interaction in APPswe cells suggesting an activation of LD degradation, which is supported by the increase of lysosome number and volume and of LD-LC3 interaction. It has been previously reported a LD accumulation in autophagy-deficient hepatocytes and MEFs [69]. The same authors also observed that LD are intimately associated with LC3-labeled autophagosomes and Lamp1-labeled lysosomes during LD catabolism by autophagy [69]. Another study performed in AML12 mouse hepatocytes showed that both p62 and LC3-labeled autophagosomes colocalize with LD under autophagy stimulating conditions [71]. Following the initial observation that small portions of larger LD can be degraded by macroautophagy after their engulfment by autophagosomes [13], we observed an accumulation of LD in cells exposed to OA plus CQ, the pharmacological inhibitor of the autophagic flux, under starvation in both cell lines. Increased lipid content has been reported in endothelial cells [72] and in the human hepatocarcinoma cell line (HepG2) treated with CQ, associated with an accumulation of LC3-II and an upregulation of p62 [73]. Our results support the involvement of macroautophagy in LD degradation under starvation upon OA treatment in both cell lines but excludes the role of this degradation pathway in the depletion of LD observed in AD cells.

The degradation of PLIN2 protein by CMA facilitates the lipolysis of neutral lipids stored into LD to generate FFA for energetic purposes [74]. In this process, PLIN2 and PLIN3 are transported in the lysosomal lumen for degradation through Lamp2A receptor located at the lysosomal membrane [75]. Cells expressing a mouse PLIN2 mutant that lacks a putative pentapeptide consensus sequence for CMA degradation (^414^SLKVQ) show decreased lipolysis and ATGL trafficking to LD, as well as a reduction of LD-autophagosome contacts and, consequently, an accumulation of larger and clustered LD [74]. In the present study, the depletion of LD during starvation, which is a potent physiological activator of CMA, is associated with an increase in PLIN2-Lamp2A co-localization. Moreover, the increase in LD number after OA treatment is linked to reduced CMA. It is interesting to note that APPswe cells present an upregulation of CMA under basal conditions, but the co-localization of PLIN2-Lamp2A does not increase under starvation. The basal increase of CMA in APPswe cells can contribute to LD lipolysis and, consequently, increased FFA levels. TAG stored in LD are converted into FFA by cytosolic lipases [68,76] or by lysosomal lipases through macrolipophagy [69]. Furthermore, our observations also suggest that macroautophagy could plays a relevant role in LD degradation in starved cells upon OA exposure, despite it is not degrade LD under normal conditions. Previous studies also show that the UPS is involved in PLIN2 degradation [55,77,78] and that proteasome impairment can contribute to PLIN2 accumulation [77]. Proteasome inhibition can also activate the autophagy-lysosome pathway [54] promoting the degradation of LD by autophagy. In our hands, the blockage of UPS decreased LD number in WT cells, suggesting that this pathway can promote the autophagic degradation of LD in this cell line. Since we did not observe an altered LD number after UPS inhibition in APPswe cells, we believe that the autophagy-lysosome pathway is impaired in these cells. Our observations also suggest that CMA can be involved in PLIN2 degradation and, consequently, promote lipolysis of neutral lipids inside LD, increasing FFA levels in APPswe cells.

In relation to LD biogenesis, we observed an increase in LD number after OA treatment in both cell lines, since this FFA is easily converted to TAG, which are stored into LD [79,80]. However, APPswe cells presented a lower number of larger LD associated with a higher concentration of TAG compared with WT cells, suggesting an impairment in LD synthesis in APPswe cells. The synthesis of neutral lipids (*e.g.*, TAG) is essential for LD biogenesis, and the increase of TAG synthesis is associated with an increased formation of LD [4,7]. Moreover, the increase of LD size and volume is dependent on TAG synthesis [81] and atypical LD fusion [82]. Seipin is also crucial for normal LD formation, being a key player in determining LD size and distribution [83]. Seipin deficiency in drosophila and yeast cells results in abnormalities in the number and morphology of LD [56,84]. Similar observations were made in SUM159 cells, where the absence of seipin led to a morphological heterogeneity of LD [56]. Accordingly, we observed a decrease in seipin levels in APPswe cells, which can contribute for the alterations in the number and morphology of LD found in these cells. Seipin forms a cage site in the ER and provides a space in the bilayer that is relatively poor in phospholipids, where the TAG lens grows [85]. Furthermore, seipin at ER-LD contacts controls LD growth by locally facilitating the continuous TAG transfer from the ER to LD, preventing their shrinkage or leakage in the opposite direction [9,11]. All these studies show that cells lacking seipin can form LD, however, they present a heterogeneous morphology and abnormal lipid and protein composition, which can result in alterations in lipid storage leading to cellular dysfunction [56]. We also found increased levels of ACC1 protein that catalyses the carboxylation of acetyl-CoA to malonyl-CoA, which is the substrate to produce long-chain saturated FA [86]. Furthermore, malonyl-CoA inhibits the interaction of FA and carnitine to prevent the transport of FA to mitochondria for oxidation and degradation [87]. The deletion of *ACC1* in A549 and H157 non-small-cell lung cancer (NSCLC) clones led to a complete loss of FA synthesis, namely palmitic, stearic, and oleic acids [88]. The increase of ACC1 protein levels was also found in the prefrontal cortex and cerebellum in a valproic acid rat model of autism [89]. Based on our observations, we hypothesized that in APPswe cells the increase in TAG synthesis in the ER bilayer promotes the formation of lens-like structures, the precursors of new LD, rich in neutral lipids. However, the downregulation of seipin levels decreases the number of places for LD growth, resulting in the formation of larger mature LD in the seipin puncta and increased TAG fluxes to the nascent LD.

Oxidative distress and damage are interlinked and clearly involved in AD pathophysiology, as demonstrated in brain, cerebrospinal fluid, and plasma from AD patients, as well as in preclinical models of the disease [90,91]. The storage of lipids into LD prevents lipotoxicity caused by lipid accumulation [92,93]. In the present study an increase in FFA and TAG was observed in APPswe cells, accompanied by increased cytosolic and mitochondrial ROS generation and lipid peroxidation. In accordance, other studies showed that high concentrations of FFA lead to lipid overload and oxidative distress in L-02 cells [94]. Moreover, the accumulation and storage of TAG in LD protect CHO and 25RA cells against FA-induced lipotoxicity [95].

Since our analyses revealed an increase in TAG levels in APPswe cells, we assessed whether TAG synthesis inhibition, using A922500, a DGAT1 pharmacological inhibitor, impact LD formation. The inhibition of DGAT1 decreased the number and size of LD in APPswe cells, which is consistent with previous reports [96–98]. The decrease in TAG synthesis leads to LD size reduction [99]. Importantly, DGAT1 inhibition also increased the close ER-mitochondria contacts and improved Δψm in APPswe cells. These results suggest that DGAT1 inhibition and, consequently, the decrease of TAG levels ameliorate APPswe cell lipid homeostasis. Since the inhibition of DGAT1 decreases TAG synthesis, it is unclear if the associated increase of ER-mitochondria contacts observed in APPswe cells represent a compensatory mechanism to promote DAG degradation (avoiding their accumulation), since several enzymes involved in DAG metabolism are MAM-resident proteins [100,101] facilitating lipid exchange between the ER and mitochondria. In addition to being metabolized into TAG, DAG can be also converted into other lipids, including FA, monoacylglycerol (MAG), phosphatidic acid (PA), which would be available for re-synthesis of phosphatidylinositol (PI) and replenishment of PIP2 (phosphatidylinositol 4,5-bisphosphate), and phosphatidylcholine (PC) or phosphatidylethanolamine (PE) by the Kennedy pathway [102]. The binding of cytidine diphosphate (CDP)-ethanolamine or CDP-choline with DAG generates PE or PC, respectively, in the ER membrane by MAM-resident proteins. Then, in the ER, PC can be converted to phosphatidylserine (PS) by the MAM-resident PS synthase 1 and imported to mitochondria, as well as PE, to be used for the synthesis of other phospholipids [42,103]. Furthermore, we observed an increase in Δψm in the presence of the DGAT1 inhibitor. However, it was previously shown a decrease of LD number and Δψm after treatment of U251 cells with A922500 [97]. The same authors reported that DGAT1 knockdown in U251 cells does not result in FFA accumulation but instead in an increase of acetyl-CoA (the major product of FA β-oxidation in mitochondria), as well as an increase of major structural lipids (PC and PE), leading to lipid dyshomeostasis and oxidative distress [97]. The differences observed between studies can be associated with distinct experimental conditions and cell types used. In the presence of the DGAT1 inhibitor, we observed a decrease in LD formation, which can promote the availability of FFA for mitochondrial β-oxidation, leading to a higher Δψm. The acetyl-CoA, NADH, and FADH_2_ are products of mitochondrial FA β-oxidation that can be transferred to the electron transport chain, generating a higher electrochemical proton gradient across the inner mitochondrial membrane causing an enhanced mitochondrial polarization [104,105].

## 5. Conclusions

Overall, our results show that neuroblastoma cells overexpressing the APPswe mutation are characterized by lipid dyshomeostasis, particularly altered LD turnover and metabolism, which affect the interaction between the ER and mitochondria and mitochondria function. Furthermore, the accumulation of FFA and TAG associated with increased ROS levels can lead to lipid peroxidation and affect the redox status of the cells. These observations highlight the importance of LD in AD pathophysiology and their potential as therapeutic targets in this neurodegenerative disease. Nevertheless, more studies are needed to support our observations and to further explore the involvement of LD in this and other neurological disorders.

## Acknowledgements

We are grateful to Armanda E. Santos (CNC—Center for Neuroscience and Cell Biology, and CIBB—Center for Innovative Biomedicine, and Laboratory of Biochemistry and Biology, Faculty of Pharmacy, University of Coimbra, Coimbra, Portugal) who kindly provided the N2A WT and N2A APPswe cell lines for this study, which were a generous gift from Ciro Isidoro (Laboratory of Molecular Pathology, Department of Health Sciences, Università del Piemonte Orientale “A. Avogadro”, Novara, Italy). We thank Mass Analytica (https://mass-analytica.com/) for providing the Lipostar 2 software, for the mass spectrometry lipid data analysis.

## Author contributions

Conceptualization: T.F., M.R.D., C.F.P., and P.I.M.; methodology: T.F., M.C., T.M, B.B.N., C.C., R.R., R.F., P.P., M.R.D., C.F.P., and P.I.M.; formal analysis: T.F. and T.M.; investigation: T.F., M.C., T.M., B.B.N., C.C., R.R., and R.F.; writing-original draft preparation: T.F.; writing-review and editing: M.R.D., C.F.P., and P.I.M.; funding acquisition: M.R.D., C.F.P., and P.I.M.; supervision: M.R.D., C.F.P., and P.I.M. All authors have read and agreed to the present version of the manuscript.

## Data Availability Statement

All data are available upon contact corresponding authors.

**Abbreviations**
Δψm: mitochondrial membrane potential
Aß: amyloid ß
ACAT/SOAT: acyl-CoA:cholesterol O-acyltransferases
ACC1: acetyl-CoA carboxylase 1
AD: Alzheimer’s disease
APP: amyloid precursor protein
APPswe: N2A cells overexpressing the human APP with the familial Swedish mutation
ATGL: adipose triglyceride lipase
BCA: bicinchoninic acid
BSA: bovine serum albumin
CE: cholesteryl ester
CMA: chaperone-mediated autophagy
CQ: chloroquine
cyt P450: cytochrome P450
DAG: diacylglycerol
DCFH2-DA: dichlorodihydrofluorescein diacetate
DGAT1/2: diacylglycerol transferase 1/2
DMEM: Dulbecco’s modified Eagle’s medium
ER: endoplasmic reticulum
FA: fatty acid
FAD: familial Alzheimer’s disease
FBS: fetal bovine serum
FFA: free fatty acid
Hsc70: heat shock-associated protein 70
IF: immunofluorescence
Lamp1 or 2A: lysosome-associated membrane receptor 1 or 2A
LC3: microtubule-associated protein light chain 3
LD: lipid droplet
MAM: mitochondria-associated membranes
MEFs: mouse embryonic fibroblasts
N2A: mouse neuroblastoma cell line
OA: oleic acid
OMM: outer mitochondrial membrane
PBS: phosphate-buffered saline
PC: phosphatidylcholine
PE: phosphatidylethanolamine
PFA: paraformaldehyde
PLA: proximity ligation assay
PLIN: perilipin
PSEN1/2: presenilin-1/2
PVDF: polyvinylidene fluoride
RIPA: radioimmunoprecipitation assay buffer
ROS: reactive oxygen species
RT: room temperature
SAD: sporadic Alzheimer’s disease
SDS-PAGE: sodium dodecyl sulfate polyacrylamide gel electrophoresis
TAG: triacylglycerol
TBA: thiobarbituric acid
TBARS: TBA reactive substracts
TBS-T: tris-buffered saline containing Tween-20
TCA: trichloroacetic acid
TF: total fraction
TMRE: tetramethylrhodamine ethyl ester perchlorate
TOM70: translocase of the mitochondrial outer membrane 70
UPS: ubiquitin-proteosome system
WB: Western blot
WT: wild-type
XIC: extracted-ion chromatogram

## References

[1] B.C. Farmer, A.E. Walsh, J.C. Kluemper, L.A. Johnson, Lipid droplets in neurodegenerative disorders, Front Neurosci 14 (2020) 742. 10.3389/fnins.2020.00742.

[2] T. Fernandes, M.R. Domingues, P.I. Moreira, C.F. Pereira, A perspective on the link between mitochondria-associated membranes (MAMs) and lipid droplets metabolism in neurodegenerative diseases, Biology (Basel) 12 (2023) 414. 10.3390/biology12030414.

[3] S. Cohen, Lipid droplets as organelles, in: Int Rev Cell Mol Biol, NIH Public Access, 2018: pp. 83–110. 10.1016/bs.ircmb.2017.12.007.

[4] C. Thiele, J. Spandl, Cell biology of lipid droplets, Curr Opin Cell Biol 20 (2008) 378–385. 10.1016/j.ceb.2008.05.009.

[5] A.R. Kimmel, C. Sztalryd, The perilipins: major cytosolic lipid droplet-associated proteins and their roles in cellular lipid storage, mobilization, and systemic homeostasis, Annu Rev Nutr 36 (2016) 471–509. 10.1146/annurev-nutr-071813-105410.

[6] M. Gao, X. Huang, B.-L. Song, H. Yang, The biogenesis of lipid droplets: lipids take center stage., Prog Lipid Res 75 (2019) 100989. 10.1016/j.plipres.2019.100989.

[7] J.A. Olzmann, P. Carvalho, Dynamics and functions of lipid droplets, Nat Rev Mol Cell Biol 20 (2019) 137–155. 10.1038/s41580-018-0085-z.

[8] T.C. Walther, R. V. Farese, Lipid droplets and cellular lipid metabolism, Annu Rev Biochem 81 (2012) 687–714. 10.1146/annurev-biochem-061009-102430.

[9] V.T. Salo, I. Belevich, S. Li, L. Karhinen, H. Vihinen, C. Vigouroux, J. Magré, C. Thiele, M. Hölttä-Vuori, E. Jokitalo, E. Ikonen, Seipin regulates ER-lipid droplet contacts and cargo delivery, EMBO J 35 (2016) 2699–2716. 10.15252/EMBJ.201695170.

[10] Y. Combot, V.T. Salo, G. Chadeuf, M. Hölttä, K. Ven, I. Pulli, S. Ducheix, C. Pecqueur, O. Renoult, B. Lak, S. Li, L. Karhinen, I. Belevich, C. Le May, J. Rieusset, S. Le Lay, M. Croyal, K.S. Tayeb, H. Vihinen, E. Jokitalo, K. Törnquist, C. Vigouroux, B. Cariou, J. Magré, A. Larhlimi, E. Ikonen, X. Prieur, Seipin localizes at endoplasmic-reticulum-mitochondria contact sites to control mitochondrial calcium import and metabolism in adipocytes, Cell Rep 38 (2022) 110213. 10.1016/j.celrep.2021.110213.

[11] V.T. Salo, S. Li, H. Vihinen, M. Hölttä-Vuori, A. Szkalisity, P. Horvath, I. Belevich, J. Peränen, C. Thiele, P. Somerharju, H. Zhao, A. Santinho, A.R. Thiam, E. Jokitalo, E. Ikonen, Seipin facilitates triglyceride flow to lipid droplet and counteracts droplet ripening via endoplasmic reticulum contact, Dev Cell 50 (2019) 478–493.e9. 10.1016/J.DEVCEL.2019.05.016.

[12] T.B. Nguyen, J.A. Olzmann, Lipid droplets and lipotoxicity during autophagy, Autophagy 13 (2017) 2002–2003. 10.1080/15548627.2017.1359451.

[13] A.D. Barbosa, S. Siniossoglou, Function of lipid droplet-organelle interactions in lipid homeostasis, Biochim Biophys Acta Mol Cell Res 1864 (2017) 1459–1468. 10.1016/j.bbamcr.2017.04.001.

[14] N. Martinez-Lopez, M. Garcia-Macia, S. Sahu, D. Athonvarangkul, E. Liebling, P. Merlo, F. Cecconi, G.J. Schwartz, R. Singh, Autophagy in the CNS and Periphery Coordinate Lipophagy and Lipolysis in the Brown Adipose Tissue and Liver, Cell Metab 23 (2016) 113–127. 10.1016/j.cmet.2015.10.008.

[15] S. Zhang, X. Peng, S. Yang, X. Li, M. Huang, S. Wei, J. Liu, G. He, H. Zheng, L. Yang, H. Li, Q. Fan, The regulation, function, and role of lipophagy, a form of selective autophagy, in metabolic disorders, Cell Death Dis 13 (2022) 132. 10.1038/s41419-022-04593-3.

[16] J. Dempsey, G. Ioannou, R. Carr, Mechanisms of Lipid Droplet Accumulation in Steatotic Liver Diseases, Semin Liver Dis 43 (2023). 10.1055/A-2186-3557.

[17] E.M. Mastoridou, A.C. Goussia, P. Kanavaros, A. V. Charchanti, Involvement of lipophagy and chaperone-mediated autophagy in the pathogenesis of non-alcoholic fatty liver disease by regulation of lipid droplets, Int J Mol Sci 24 (2023) 15891. 10.3390/ijms242115891.

[18] X. Guo, Q. Shi, W. Zhang, Z. Qi, H. Lv, F. Man, Y. Xie, Y. Zhu, J. Zhang, Lipid droplet-a new target in ischemic heart disease, J Cardiovasc Transl Res 15 (2022) 730–739. 10.1007/S12265-021-10204-X.

[19] D. Yang, X. Wang, L. Zhang, Y. Fang, Q. Zheng, X. Liu, W. Yu, S. Chen, J. Ying, F. Hua, Lipid metabolism and storage in neuroglia: role in brain development and neurodegenerative diseases, Cell Biosci 12 (2022) 106. 10.1186/s13578-022-00828-0.

[20] Y. Liu, A. Thalamuthu, K.A. Mather, J. Crawford, M. Ulanova, M.W.K. Wong, R. Pickford, P.S. Sachdev, N. Braidy, Plasma lipidome is dysregulated in Alzheimer’s disease and is associated with disease risk genes, Transl Psychiatry 11 (2021) 344. 10.1038/s41398-021-01362-2.

[21] V.R. Varma, H. Büşra Lüleci, A.M. Oommen, S. Varma, C.T. Blackshear, M.E. Griswold, Y. An, J.A. Roberts, R. O’Brien, O. Pletnikova, J.C. Troncoso, D.A. Bennett, T. Çakır, C. Legido-Quigley, M. Thambisetty, Abnormal brain cholesterol homeostasis in Alzheimer’s disease—a targeted metabolomic and transcriptomic study, NPJ Aging Mech Dis 7 (2021) 11. 10.1038/s41514-021-00064-9.

[22] F. Dakterzada, M. Jové, R. Huerto, A. Carnes, J. Sol, R. Pamplona, G. Piñol-Ripoll, Cerebrospinal fluid neutral lipids predict progression from mild cognitive impairment to Alzheimer’s disease, Geroscience 46 (2023) 683–696. 10.1007/s11357-023-00989-x.

[23] P.T. Nelson, E.L. Abner, F.A. Schmitt, R.J. Kryscio, G.A. Jicha, K. Santacruz, C.D. Smith, E. Patel, W.R. Markesbery, Brains with medial temporal lobe neurofibrillary tangles but no neuritic amyloid plaques are a diagnostic dilemma but may have pathogenetic aspects distinct from Alzheimer disease, J Neuropathol Exp Neurol 68 (2009) 774–784. 10.1097/NEN.0b013e3181aacbe9.

[24] B. Winblad, P. Amouyel, S. Andrieu, C. Ballard, C. Brayne, H. Brodaty, A. Cedazo-Minguez, B. Dubois, D. Edvardsson, H. Feldman, L. Fratiglioni, G.B. Frisoni, S. Gauthier, J. Georges, C. Graff, K. Iqbal, F. Jessen, G. Johansson, L. Jönsson, M. Kivipelto, M. Knapp, F. Mangialasche, R. Melis, A. Nordberg, M.O. Rikkert, C. Qiu, T.P. Sakmar, P. Scheltens, L.S. Schneider, R. Sperling, L.O. Tjernberg, G. Waldemar, A. Wimo, H. Zetterberg, Defeating Alzheimer’s disease and other dementias: a priority for European science and society, Lancet Neurol 15 (2016) 455–532. 10.1016/S1474-4422(16)00062-4.

[25] T. Fernandes, R. Resende, D.F. Silva, A.P. Marques, A.E. Santos, S.M. Cardoso, M.R. Domingues, P.I. Moreira, C.F. Pereira, Structural and functional alterations in mitochondria-associated membranes (MAMs) and in mitochondria activate stress response mechanisms in an in vitro model of Alzheimer’s disease, Biomedicines 9 (2021) 881. 10.3390/biomedicines9080881.

[26] E. Area-Gomez, A. de Groof, E. Bonilla, J. Montesinos, K. Tanji, I. Boldogh, L. Pon, E.A. Schon, A key role for MAM in mediating mitochondrial dysfunction in Alzheimer disease, Cell Death Dis 9 (2018) 335. 10.1038/s41419-017-0215-0.

[27] E. Ferreiro, I. Baldeiras, I.L. Ferreira, R.O. Costa, A.C. Rego, C.F. Pereira, C.R. Oliveira, Mitochondrial- and endoplasmic reticulum-associated oxidative stress in Alzheimer’s disease: from pathogenesis to biomarkers, Int J Cell Biol 2012 (2012) 1–23. 10.1155/2012/735206.

[28] E. Area-Gomez, M. del Carmen Lara Castillo, M.D. Tambini, C. Guardia-Laguarta, A.J.C. de Groof, M. Madra, J. Ikenouchi, M. Umeda, T.D. Bird, S.L. Sturley, E.A. Schon, Upregulated function of mitochondria-associated ER membranes in Alzheimer disease, EMBO J 31 (2012) 4106–4123. 10.1038/emboj.2012.202.

[29] L. Liu, K.R. MacKenzie, N. Putluri, M. Maletić-Savatić, H.J. Bellen, The glia-neuron lactate shuttle and elevated ROS promote lipid synthesis in neurons and lipid droplet accumulation in glia via APOE/D, Cell Metab 26 (2017) 719–737.e6. 10.1016/j.cmet.2017.08.024.

[30] P. Wood, A. Phillipps, R.L. Woltjer, J. Kaye, J. Quinn, Increased lysophosphatidylethanolamine and diacylglycerol levels in Alzheimer’s disease plasma, JSM Alzheimers Dis Relat Dement 1 (2014).

[31] R. González-Domínguez, T. García-Barrera, J.L. Gómez-Ariza, Application of a novel metabolomic approach based on atmospheric pressure photoionization mass spectrometry using flow injection analysis for the study of Alzheimer׳s disease, Talanta 131 (2015) 480–489. 10.1016/j.talanta.2014.07.075.

[32] R.B. Chan, T.G. Oliveira, E.P. Cortes, L.S. Honig, K.E. Duff, S.A. Small, M.R. Wenk, G. Shui, G. Di Paolo, Comparative lipidomic analysis of mouse and Human brain with Alzheimer disease, Journal of Biological Chemistry 287 (2012) 2678–2688. 10.1074/jbc.M111.274142.

[33] Y. Tajima, M. Ishikawa, K. Maekawa, M. Murayama, Y. Senoo, T. Nishimaki-Mogami, H. Nakanishi, K. Ikeda, M. Arita, R. Taguchi, A. Okuno, R. Mikawa, S. Niida, O. Takikawa, Y. Saito, Lipidomic analysis of brain tissues and plasma in a mouse model expressing mutated human amyloid precursor protein/tau for Alzheimer’s disease, Lipids Health Dis 12 (2013) 68. 10.1186/1476-511X-12-68.

[34] M.S. Haney, R. Pálovics, C.N. Munson, C. Long, P.K. Johansson, O. Yip, W. Dong, E. Rawat, E. West, J.C.M. Schlachetzki, A. Tsai, I.H. Guldner, B.S. Lamichhane, A. Smith, N. Schaum, K. Calcuttawala, A. Shin, Y.-H. Wang, C. Wang, N. Koutsodendris, G.E. Serrano, T.G. Beach, E.M. Reiman, C.K. Glass, M. Abu-Remaileh, A. Enejder, Y. Huang, T. Wyss-Coray, APOE4/4 is linked to damaging lipid droplets in Alzheimer’s disease microglia, Nature 628 (2024) 154–161. 10.1038/s41586-024-07185-7.

[35] M.J. Moulton, S. Barish, I. Ralhan, J. Chang, L.D. Goodman, J.G. Harland, P.C. Marcogliese, J.O. Johansson, M.S. Ioannou, H.J. Bellen, Neuronal ROS-induced glial lipid droplet formation is altered by loss of Alzheimer’s disease–associated genes, Proceedings of the National Academy of Sciences 118 (2021). 10.1073/pnas.2112095118.

[36] G. Thinakaran, D.B. Teplow, R. Siman, B. Greenberg, S.S. Sisodia, Metabolism of the “Swedish” amyloid precursor protein variant in Neuro2a (N2a) cells, Journal of Biological Chemistry 271 (1996) 9390–9397. 10.1074/jbc.271.16.9390.

[37] R. Resende, M. Ferreira-Marques, P. Moreira, J.R.M. Coimbra, S.J. Baptista, C. Isidoro, J.A.R. Salvador, T.C.P. Dinis, C.F. Pereira, A.E. Santos, New BACE1 chimeric peptide inhibitors selectively prevent AβPP-β cleavage decreasing amyloid-β production and accumulation in Alzheimer’s disease models, Journal of Alzheimer’s Disease 76 (2020) 1317–1337. 10.3233/JAD-200381.

[38] S. Nakajima, M. Gotoh, K. Fukasawa, K. Murakami-Murofushi, H. Kunugi, Oleic acid is a potent inducer for lipid droplet accumulation through its esterification to glycerol by diacylglycerol acyltransferase in primary cortical astrocytes., Brain Res 1725 (2019) 146484. 10.1016/j.brainres.2019.146484.

[39] P. García Casas, M. Rossini, L. Påvénius, M. Saeed, N. Arnst, S. Sonda, T. Fernandes, I. D’Arsiè, M. Bruzzone, V. Berno, A. Raimondi, M.L. Sassano, L. Naia, E. Barbieri, S. Sigismund, P. Agostinis, M. Sturlese, B.A. Niemeyer, H. Brismar, M. Ankarcrona, A. Gautier, P. Pizzo, R. Filadi, Simultaneous detection of membrane contact dynamics and associated Ca2+ signals by reversible chemogenetic reporters, Nat Commun 15 (2024) 9775. 10.1038/s41467-024-52985-0.

[40] B.C. Dickinson, V.S. Lin, C.J. Chang, Preparation and use of MitoPY1 for imaging hydrogen peroxide in mitochondria of live cells, Nat Protoc 8 (2013) 1249–1259. 10.1038/nprot.2013.064.

[41] C. Carvalho, S.C. Correia, R. Seiça, P.I. Moreira, WWOX inhibition by Zfra1-31 restores mitochondrial homeostasis and viability of neuronal cells exposed to high glucose., Cell Mol Life Sci 79 (2022) 487. 10.1007/s00018-022-04508-7.

[42] T. Fernandes, T. Melo, T. Conde, B. Neves, P. Domingues, R. Resende, C.F. Pereira, P.I. Moreira, M.R. Domingues, Mapping the lipidome in mitochondria-associated membranes (MAMs) in an in vitro model of Alzheimer’s disease, J Neurochem 168 (2024) 1237– 1253. 10.1111/jnc.16072.

[43] S. Colombo, T. Melo, M. Martínez-López, M.J. Carrasco, M.R. Domingues, D. Pérez-Sala, P. Domingues, Phospholipidome of endothelial cells shows a different adaptation response upon oxidative, glycative and lipoxidative stress, Sci Rep 8 (2018) 12365. 10.1038/s41598-018-30695-0.

[44] H.B. Ferreira, T. Melo, H. Rocha, A. Paiva, P. Domingues, M.R. Domingues, Lipid profile variability in children at different ages measured in dried blood spots, Mol Omics 19 (2023) 229–237. 10.1039/D2MO00206J.

[45] C.P. Faria, B. Ferreira, Á. Lourenço, I. Guerra, T. Melo, P. Domingues, M. do R.M. Domingues, M.T. Cruz, M. do C. Sousa, Lipidome of extracellular vesicles from Giardia lamblia, PLoS One 18 (2023) e0291292. 10.1371/journal.pone.0291292.

[46] I.M.S. Guerra, H.B. Ferreira, T. Maurício, M. Pinho, L. Diogo, S. Moreira, L. Goracci, S. Bonciarelli, T. Melo, P. Domingues, M.R. Domingues, A.S.P. Moreira, Plasma lipidomics analysis reveals altered profile of triglycerides and phospholipids in children with medium-chain acyl-CoA dehydrogenase deficiency, J Inherit Metab Dis 47 (2024) 731– 745. 10.1002/jimd.12718.

[47] L. Goracci, S. Tortorella, P. Tiberi, R.M. Pellegrino, A. Di Veroli, A. Valeri, G. Cruciani, Lipostar, a comprehensive platform-neutral cheminformatics tool for lipidomics, Anal Chem 89 (2017) 6257–6264. 10.1021/acs.analchem.7b01259.

[48] M. Lange, G. Angelidou, Z. Ni, A. Criscuolo, J. Schiller, M. Blüher, M. Fedorova, AdipoAtlas: a reference lipidome for human white adipose tissue, Cell Rep Med 2 (2021) 100407. 10.1016/j.xcrm.2021.100407.

[49] S. Li, C.-W. Lu, E.C. Diem, W. Li, M. Guderian, M. Lindenberg, F. Kruse, M. Buettner, S. Floess, M.R. Winny, R. Geffers, H.-H. Richnow, W.-R. Abraham, G.A. Grassl, M. Lochner, Acetyl-CoA-Carboxylase 1-mediated de novo fatty acid synthesis sustains Lgr5+ intestinal stem cell function, Nat Commun 13 (2022) 3998. 10.1038/s41467-022-31725-2.

[50] T.C. Harned, R. V. Stan, Z. Cao, R. Chakrabarti, H.N. Higgs, C.C.Y. Chang, T.Y. Chang, Acute ACAT1/SOAT1 blockade increases MAM cholesterol and strengthens ER-mitochondria connectivity, Int J Mol Sci 24 (2023) 5525. 10.3390/ijms24065525.

[51] L. Cui, P. Liu, Two types of contact between lipid droplets and mitochondria, Front Cell Dev Biol 8 (2020) 618322. 10.3389/fcell.2020.618322.

[52] M. Kitada, D. Koya, Autophagy in metabolic disease and ageing, Nat Rev Endocrinol 17 (2021) 647–661. 10.1038/S41574-021-00551-9.

[53] M. Mauthe, I. Orhon, C. Rocchi, X. Zhou, M. Luhr, K.J. Hijlkema, R.P. Coppes, N. Engedal, M. Mari, F. Reggiori, Chloroquine inhibits autophagic flux by decreasing autophagosome-lysosome fusion, Autophagy 14 (2018) 1435–1455. 10.1080/15548627.2018.1474314.

[54] C. Li, X. Wang, X. Li, K. Qiu, F. Jiao, Y. Liu, Q. Kong, Y. Liu, Y. Wu, Proteasome inhibition activates autophagy-lysosome pathway associated with TFEB dephosphorylation and nuclear translocation, Front Cell Dev Biol 7 (2019). 10.3389/fcell.2019.00170.

[55] K.T. Nguyen, C.-S. Lee, S.-H. Mun, N.T. Truong, S.K. Park, C.-S. Hwang, N-terminal acetylation and the N-end rule pathway control degradation of the lipid droplet protein PLIN2, Journal of Biological Chemistry 294 (2019) 379–388. 10.1074/jbc.RA118.005556.

[56] H. Wang, M. Becuwe, B.E. Housden, C. Chitraju, A.J. Porras, M.M. Graham, X.N. Liu, A.R. Thiam, D.B. Savage, A.K. Agarwal, A. Garg, M.-J. Olarte, Q. Lin, F. Fröhlich, H.K. Hannibal-Bach, S. Upadhyayula, N. Perrimon, T. Kirchhausen, C.S. Ejsing, T.C. Walther, R. V Farese, Seipin is required for converting nascent to mature lipid droplets, Elife 5 (2016). 10.7554/eLife.16582.

[57] L.-J. Su, J.-H. Zhang, H. Gomez, R. Murugan, X. Hong, D. Xu, F. Jiang, Z.-Y. Peng, Reactive oxygen species-induced lipid peroxidation in apoptosis, autophagy, and ferroptosis, Oxid Med Cell Longev 2019 (2019) 1–13. 10.1155/2019/5080843.

[58] T. Fernandes, M.R. Domingues, C.F. Pereira, P.I. Moreira, Mitochondria-associated membranes (MAMs) as modulators of oxidative stress in Alzheimer disease, in: Modulation of Oxidative Stress, Elsevier, 2023: pp. 81–96. 10.1016/B978-0-443-19247-0.00012-6.

[59] C. Chitraju, N. Mejhert, J.T. Haas, L.G. Diaz-Ramirez, C.A. Grueter, J.E. Imbriglio, S. Pinto, S.K. Koliwad, T.C. Walther, R. V. Farese, Triglyceride synthesis by DGAT1 protects adipocytes from lipid-induced ER stress during lipolysis, Cell Metab 26 (2017) 407–418.e3. 10.1016/j.cmet.2017.07.012.

[60] D. Casares, P. V. Escribá, C.A. Rosselló, Membrane lipid composition: effect on membrane and organelle structure, function and compartmentalization and therapeutic avenues, Int J Mol Sci 20 (2019) 2167. 10.3390/ijms20092167.

[61] P. Campomanes, V. Zoni, S. Vanni, Local accumulation of diacylglycerol alters membrane properties nonlinearly due to its transbilayer activity, Commun Chem 2 (2019) 72. 10.1038/s42004-019-0175-7.

[62] D. Li, S.-G. Yang, C.-W. He, Z.-T. Zhang, Y. Liang, H. Li, J. Zhu, X. Su, Q. Gong, Z. Xie, Excess diacylglycerol at the endoplasmic reticulum disrupts endomembrane homeostasis and autophagy, BMC Biol 18 (2020) 107. 10.1186/s12915-020-00837-w.

[63] M.B. Khawar, H. Gao, W. Li, Autophagy and Lipid Metabolism, Adv Exp Med Biol 1206 (2019) 359–374. 10.1007/978-981-15-0602-4_17.

[64] H. Chew, V.A. Solomon, A.N. Fonteh, Involvement of lipids in Alzheimer’s disease pathology and potential therapies, Front Physiol 11 (2020) 1–28. 10.3389/fphys.2020.00598.

[65] M.W. Wong, N. Braidy, A. Poljak, R. Pickford, M. Thambisetty, P.S. Sachdev, Dysregulation of lipids in Alzheimer’s disease and their role as potential biomarkers, Alzheimer’s & Dementia 13 (2017) 810–827. 10.1016/j.jalz.2017.01.008.

[66] L.K. Hamilton, M. Dufresne, S.E. Joppé, S. Petryszyn, A. Aumont, F. Calon, F. Barnabé-Heider, A. Furtos, M. Parent, P. Chaurand, K.J.L. Fernandes, Aberrant lipid metabolism in the forebrain niche suppresses adult neural stem cell proliferation in an animal model of Alzheimer’s disease, Cell Stem Cell 17 (2015) 397–411. 10.1016/j.stem.2015.08.001.

[67] B. Hutter-Paier, H.J. Huttunen, L. Puglielli, C.B. Eckman, D.Y. Kim, A. Hofmeister, R.D. Moir, S.B. Domnitz, M.P. Frosch, M. Windisch, D.M. Kovacs, The ACAT inhibitor CP-113,818 markedly reduces amyloid pathology in a mouse model of Alzheimer’s disease., Neuron 44 (2004) 227–38. 10.1016/j.neuron.2004.08.043.

[68] A.D. Quiroga, R. Lehner, Liver triacylglycerol lipases, Biochimica et Biophysica Acta (BBA) - Molecular and Cell Biology of Lipids 1821 (2012) 762–769. 10.1016/j.bbalip.2011.09.007.

[69] R. Singh, S. Kaushik, Y. Wang, Y. Xiang, I. Novak, M. Komatsu, K. Tanaka, A.M. Cuervo, M.J. Czaja, Autophagy regulates lipid metabolism, Nature 458 (2009) 1131– 1135. 10.1038/nature07976.

[70] R. Zechner, F. Madeo, D. Kratky, Cytosolic lipolysis and lipophagy: two sides of the same coin, Nat Rev Mol Cell Biol 18 (2017) 671–684. 10.1038/nrm.2017.76.

[71] L. Wang, J. Zhou, S. Yan, G. Lei, C.-H. Lee, X.-M. Yin, Ethanol-triggered lipophagy requires SQSTM1 in AML12 hepatic cells, Sci Rep 7 (2017) 12307. 10.1038/s41598-017-12485-2.

[72] E. Bik, L. Mateuszuk, J. Orleanska, M. Baranska, S. Chlopicki, K. Majzner, Chloroquine-induced accumulation of autophagosomes and lipids in the endothelium, Int J Mol Sci 22 (2021) 1–14. 10.3390/IJMS22052401.

[73] F. Xu, H.M. Tautenhahn, O. Dirsch, U. Dahmen, Blocking autophagy with chloroquine aggravates lipid accumulation and reduces intracellular energy synthesis in hepatocellular carcinoma cells, both contributing to its anti-proliferative effect, J Cancer Res Clin Oncol 148 (2022) 3243. 10.1007/S00432-022-04074-2.

[74] S. Kaushik, A.M. Cuervo, Degradation of lipid droplet-associated proteins by chaperone-mediated autophagy facilitates lipolysis, Nat Cell Biol 17 (2015) 759–770. 10.1038/NCB3166.

[75] K. Drizyte-Miller, M.B. Schott, M.A. McNiven, Lipid droplet contacts with autophagosomes, lysosomes, and other degradative vesicles, Contact 3 (2020) 251525642091089. 10.1177/2515256420910892.

[76] A. Lass, R. Zimmermann, M. Oberer, R. Zechner, Lipolysis – a highly regulated multi-enzyme complex mediates the catabolism of cellular fat stores, Prog Lipid Res 50 (2011) 14–27. 10.1016/j.plipres.2010.10.004.

[77] E. Pisano, L. Pacifico, F.M. Perla, G. Liuzzo, C. Chiesa, M. Lavorato, G. Mingrone, M. Fabrizi, D. Fintini, A. Severino, M. Manco, Upregulated monocyte expression of PLIN2 is associated with early arterial injury in children with overweight/obesity, Atherosclerosis 327 (2021) 68–75. 10.1016/j.atherosclerosis.2021.04.016.

[78] Y. Masuda, H. Itabe, M. Odaki, K. Hama, Y. Fujimoto, M. Mori, N. Sasabe, J. Aoki, H. Arai, T. Takano, ADRP/adipophilin is degraded through the proteasome-dependent pathway during regression of lipid-storing cells, J Lipid Res 47 (2006) 87–98. 10.1194/jlr.M500170-JLR200.

[79] D.F. Dibwe, E. Kitayama, S. Oba, N. Takeishi, H. Chiba, S.-P. Hui, Inhibition of lipid accumulation and oxidation in hepatocytes by bioactive bean extracts, Antioxidants 13 (2024) 513. 10.3390/antiox13050513.

[80] A. Rohwedder, Q. Zhang, S.A. Rudge, M.J.O. Wakelam, Lipid droplet formation in response to oleic acid in Huh-7 cells is a fatty acid receptor mediated event, J Cell Sci 127 (2014) 3104–3115. 10.1242/jcs.145854.

[81] D.J. Murphy, The dynamic roles of intracellular lipid droplets: from archaea to mammals, Protoplasma 249 (2012) 541–585. 10.1007/s00709-011-0329-7.

[82] W. Xu, L. Wu, M. Yu, F.-J. Chen, M. Arshad, X. Xia, H. Ren, J. Yu, L. Xu, D. Xu, J.Z. Li, P. Li, L. Zhou, Differential roles of cell death-inducing DNA fragmentation factor-α-like effector (CIDE) proteins in promoting lipid droplet fusion and growth in subpopulations of hepatocytes, Journal of Biological Chemistry 291 (2016) 4282–4293. 10.1074/jbc.M115.701094.

[83] W. Fei, X. Du, H. Yang, Seipin, adipogenesis and lipid droplets, Trends in Endocrinology & Metabolism 22 (2011) 204–210. 10.1016/j.tem.2011.02.004.

[84] B.R. Cartwright, D.D. Binns, C.L. Hilton, S. Han, Q. Gao, J.M. Goodman, Seipin performs dissectible functions in promoting lipid droplet biogenesis and regulating droplet morphology, Mol Biol Cell 26 (2015) 726–739. 10.1091/mbc.E14-08-1303.

[85] H. Arlt, X. Sui, B. Folger, C. Adams, X. Chen, R. Remme, F.A. Hamprecht, F. DiMaio, M. Liao, J.M. Goodman, R. V. Farese, T.C. Walther, Seipin forms a flexible cage at lipid droplet formation sites, Nat Struct Mol Biol 29 (2022) 194–202. 10.1038/s41594-021-00718-y.

[86] Y. Wang, W. Yu, S. Li, D. Guo, J. He, Y. Wang, Acetyl-CoA carboxylases and diseases, Front Oncol 12 (2022) 836058. 10.3389/fonc.2022.836058.

[87] D.W. Foster, Malonyl-CoA: the regulator of fatty acid synthesis and oxidation, Journal of Clinical Investigation 122 (2012) 1958–1959. 10.1172/JCI63967.

[88] R.U. Svensson, S.J. Parker, L.J. Eichner, M.J. Kolar, M. Wallace, S.N. Brun, P.S. Lombardo, J.L. Van Nostrand, A. Hutchins, L. Vera, L. Gerken, J. Greenwood, S. Bhat, G. Harriman, W.F. Westlin, H.J. Harwood, A. Saghatelian, R. Kapeller, C.M. Metallo, R.J. Shaw, Inhibition of acetyl-CoA carboxylase suppresses fatty acid synthesis and tumor growth of non-small-cell lung cancer in preclinical models, Nat Med 22 (2016) 1108– 1119. 10.1038/nm.4181.

[89] J. Chen, W. Wu, Y. Fu, S. Yu, D. Cui, M. Zhao, Y. Du, J. Li, X. Li, Increased expression of fatty acid synthase and acetyl-CoA carboxylase in the prefrontal cortex and cerebellum in the valproic acid model of autism, Exp Ther Med 12 (2016) 1293–1298. 10.3892/etm.2016.3508.

[90] L. Ferré-González, C. Peña-Bautista, M. Baquero, C. Cháfer-Pericás, Assessment of lipid peroxidation in Alzheimer’s disease differential diagnosis and prognosis, Antioxidants 11 (2022) 551. 10.3390/antiox11030551.

[91] S. Arlt, U. Beisiegel, A. Kontush, Lipid peroxidation in neurodegeneration: new insights into Alzheimer’s disease, Curr Opin Lipidol 13 (2002) 289–294. 10.1097/00041433-200206000-00009.

[92] J.C. Arroyave-Ospina, Z. Wu, Y. Geng, H. Moshage, Role of oxidative stress in the pathogenesis of non-alcoholic fatty liver disease: implications for prevention and therapy, Antioxidants 10 (2021) 174. 10.3390/antiox10020174.

[93] P. Xie, J.G. Zhu, L.X. Wang, Y. Liu, E.J. Diao, D.Q. Gong, T.W. Liu, Lipid accumulation and oxidative stress in the crop tissues of male and female pigeons during incubation and chick-rearing periods, Poult Sci 102 (2023) 102289. 10.1016/j.psj.2022.102289.

[94] S. Qin, J. Yin, K. Huang, Free Fatty Acids Increase Intracellular Lipid Accumulation and Oxidative Stress by Modulating PPARα and SREBP-1c in L-02 Cells, Lipids 51 (2016) 797–805. 10.1007/S11745-016-4160-Y.

[95] L.L. Listenberger, X. Han, S.E. Lewis, S. Cases, R. V Farese, D.S. Ory, J.E. Schaffer, Triglyceride accumulation protects against fatty acid-induced lipotoxicity, Proc Natl Acad Sci U S A 100 (2003) 3077–82. 10.1073/pnas.0630588100.

[96] A. Kuo, M.Y. Lee, W.C. Sessa, Lipid droplet biogenesis and function in the endothelium, Circ Res 120 (2017) 1289–1297. 10.1161/CIRCRESAHA.116.310498.

[97] X. Cheng, F. Geng, M. Pan, X. Wu, Y. Zhong, C. Wang, Z. Tian, C. Cheng, R. Zhang, V. Puduvalli, C. Horbinski, X. Mo, X. Han, A. Chakravarti, D. Guo, Targeting DGAT1 ameliorates glioblastoma by increasing fat catabolism and oxidative stress, Cell Metab 32 (2020) 229–242.e8. 10.1016/j.cmet.2020.06.002.

[98] L. Teixeira, F.S. Pereira-Dutra, P.A. Reis, T. Cunha-Fernandes, M.Y. Yoshinaga, L. Souza-Moreira, E.K. Souza, E.A. Barreto, T.P. Silva, H. Espinheira-Silva, T. Igreja, M.M. Antunes, A.C.S. Bombaça, C.F. Gonçalves-de-Albuquerque, G.B. Menezes, E.D. Hottz, R.F.S. Menna-Barreto, C.M. Maya-Monteiro, F.A. Bozza, S. Miyamoto, R.C.N. Melo, P.T. Bozza, Prevention of lipid droplet accumulation by DGAT1 inhibition ameliorates sepsis-induced liver injury and inflammation, JHEP Reports 6 (2024) 100984. 10.1016/j.jhepr.2023.100984.

[99] Y.-S. Chen, H.-M. Liu, T.-Y. Lee, Ursodeoxycholic acid regulates hepatic energy homeostasis and white adipose tissue macrophages polarization in leptin-deficiency obese mice, Cells 8 (2019) 253. 10.3390/cells8030253.

[100] S.J. Stone, J.E. Vance, Phosphatidylserine synthase-1 and -2 are localized to mitochondria-associated membranes, Journal of Biological Chemistry 275 (2000) 34534–34540. 10.1074/jbc.M002865200.

[101] Z. Cui, J.E. Vance, M.H. Chen, D.R. Voelker, D.E. Vance, Cloning and expression of a novel phosphatidylethanolamine N-methyltransferase. A specific biochemical and cytological marker for a unique membrane fraction in rat liver, Journal of Biological Chemistry 268 (1993) 16655–16663. 10.1016/S0021-9258(19)85468-6.

[102] T.O. Eichmann, A. Lass, DAG tales: the multiple faces of diacylglycerol— stereochemistry, metabolism, and signaling, Cellular and Molecular Life Sciences 72 (2015) 3931–3952. 10.1007/s00018-015-1982-3.

[103] I. Gordaliza-Alaguero, C. Cantó, A. Zorzano, Metabolic implications of organelle– mitochondria communication, EMBO Rep 20 (2019) 1–27. 10.15252/embr.201947928.

[104] M.M. Adeva-Andany, N. Carneiro-Freire, M. Seco-Filgueira, C. Fernández-Fernández, D. Mouriño-Bayolo, Mitochondrial β-oxidation of saturated fatty acids in humans, Mitochondrion 46 (2019) 73–90. 10.1016/j.mito.2018.02.009.

[105] H.M. Begum, K. Shen, Intracellular and microenvironmental regulation of mitochondrial membrane potential in cancer cells, WIREs Mechanisms of Disease 15 (2023). 10.1002/wsbm.1595.

